# Oncogene-Mechanics Axis: KRAS G12C Confers Agility Enabling Malignant Mechano-responses to Peristalsis in Colorectal Cancer

**DOI:** 10.64898/2026.04.28.721453

**Authors:** Astha Lamichhane, Vsevolod Cheburkanov, Mykyta Kizilov, Abhinav Shenoy, Audrey G. Head, Vladislav Yakovlev, Shreya A. Raghavan

## Abstract

Oncogene activity and mechanical forces individually and collectively drive colorectal cancer, yet the integration of these signals is unknown. We used a patented peristalsis bioreactor to determine how oncogenic KRAS G12C mutations alter the cellular response to colonic peristalsis. Although both healthy intestinal cells (HIEC-6) and KRAS G12C cancer cells (SW837) sensed peristalsis via ERK phosphorylation (1.7-4.7X increase), their mechano-responses diverged significantly. Peristalsis triggered a 9-fold enrichment of LGR5^+^ cancer stem cells in KRAS G12C cancer cells, an effect absent in healthy HIEC-6 controls. Using Brillouin microscopy, we discovered that KRAS G12C possessed a mechanically primed state exhibiting 9% reduction in Brillouin Full Width at Half Maximum, indicative of lower intracellular viscosity. Peristalsis further decreased intracellular viscosity in KRAS G12C cells by 11.9%, accompanied by pronounced cell elongation, whereas HIEC-6 maintained viscoelastic homeostasis. This KRAS-dependent mechanical agility enabled KRAS G12C cancer cells to leverage, rather than resist peristalsis, resulting in LGR5 enrichment and malignant progression. Pharmacologic KRAS G12C inhibition with sotorasib reduced LGR5^+^ cells by 50% under peristalsis, reversing the malignant mechanotype. Conversely, introducing KRAS G12C into HIEC-6 disrupted their mechanical homeostasis and significantly increased LGR5, recapitulating the KRAS-dependent mechano-phenotype. Our findings identify a novel KRAS oncogene-mechanics axis, suggesting that targeting the cell’s mechanical state could be a powerful complement to emerging KRAS-directed therapies.

**Teaser:** How does the gut’s movement fuel cancer? We report a novel oncogenic KRAS-mechanics axis, where KRAS G12C mutations lower cancer cell viscosity, conferring agility and deformability. This allows cancer cells to leverage colonic peristalsis resulting in cancer stem cell enrichment. Cells transform physical forces associated with peristalsis into a powerful driver of tumor progression.

## Introduction

Colorectal cancer (CRC) is a leading cause of cancer mortality globally^1^. Among the most prevalent genetic drivers are mutations in the Kirsten rat sarcoma virus (*KRAS*) gene, which occur in ∼40% of CRC cases. Oncogenic mutations in *KRAS* promote uncontrolled growth, survival and aberrant cytoskeletal remodeling through persistent activation of downstream effector pathways^2^. Within this mutational landscape, *KRAS* G12C stands out as a distinct allele, linked to poor therapeutic response and aggressive tumor behavior^3^. Interestingly, the G12C mutation is also selectively targetable by covalent inhibitors^4^, positioning this variant as a unique intersection of aggressive tumor biology and druggable vulnerability in a subset of colorectal cancer patients^5^. Here, we expose the vulnerability of *KRAS* G12C mutant colorectal cancer related to the unique colorectal tumor microenvironment.

Beyond cell intrinsic oncogene signaling, mechanics of the tumor microenvironment drive colorectal cancer progression^6^. The pattern of mechanics is unique in the colon tumor microenvironment which is subject to rhythmic deformation due to colonic peristalsis^7^. We invented a peristalsis bioreactor device capable of externally applying peristalsis mechanics to colorectal cancer cell cultures in vitro^8,9^. Using the bioreactor’s single platform rigor, our prior work also strongly demonstrated that peristalsis was a unique mechanical stimulus, eliciting biological mechano-sensitive responses different from fluid shear stress or cyclic strain alone^9–11^. Sensing bioreactor-applied peristalsis, *KRAS* mutant colorectal cancer cells enriched LGR5 cancer stem cells, leading to increased tumorigenicity^10,11^. In stark contrast, colorectal cancer cells that lacked *KRAS* oncogene activation failed to activate stemness under peristalsis, suggesting that oncogenic KRAS may alter how cells sense and transduce mechanics. LGR5 enrichment is a central, and clinically relevant readout for malignant progression in colorectal cancer. Clinically, LGR5 is overexpressed in colorectal cancer tissues compared to matched healthy epithelium, with its elevation beginning early during the adenoma to carcinoma transition and continuing to rise during malignant progression leading to invasion and metastasis^12^. This makes LGR5 enrichment an established and essential readout downstream of mechanical stimulation in our current study.

In response to mechanics, cells change their biochemical programming to enable functional outcomes like deformation and migration^10,13^. How a cell responds to mechanics, however, is fundamentally governed by its intrinsic material properties like viscosity and elasticity. Together, these properties impact a cell’s ability to deform in response to mechanics. Oncogenic transformation changes cellular material properties like stiffness and viscoelasticity compared to healthy counterparts, occurring partly via altered cytoskeletal architecture ^13–17^. Yet in CRC, the understanding of cellular material and mechanical properties is still emerging, especially as it pertains to *KRAS* oncogene activation.

In the current study, we hypothesized that oncogenic *KRAS* promotes a malignant mechano-response to peristalsis first by altering cellular material and mechanical properties. To investigate, we used Brillouin microscopy, a powerful label-free optical modality for non-invasive mapping of mechanical properties in biological systems with sub-cellular resolution ^18,19^. Brillouin microscopy relies on inelastic scattering of light from thermally driven density fluctuations, providing direct access to high-frequency viscoelastic response. This inelastic scattering event yields two measurable parameters: the scattered photon frequency shift (*ν_B_*, Brillouin shift) and the Brillouin linewidth (*Γ_B_*, full width at half maximum, FWHM). The squared Brillouin shift (*ν_B_*^2^) is proportional to the material’s high-frequency elastic modulus, while the product of the Brillouin shift and linewidth (*ν_B_Γ_B_*) is proportional to its high-frequency viscosity. Together, these relationships enable quantitative assessment and comparison of cellular viscoelastic properties under different experimental conditions. Recent advances ^20,21^ have established Brillouin microscopy as a robust platform for biological imaging, enabling quantitative, spatially resolved measurements of viscoelastic properties at the microscale. Its noninvasive nature, minimal sample preparation, and intrinsic ability to simultaneously capture both elastic and viscous responses distinguish it from contact-based techniques such as atomic force microscopy, positioning Brillouin microscopy as a versatile tool for probing the physical underpinnings of biological function.

Combining Brillouin microscopy with molecular and functional analyses, our studies demonstrate that oncogenic *KRAS* G12C established a distinct intracellular mechanical phenotype characterized by reduced intracellular viscosity. Peristalsis takes advantage of this agile mechano-phenotype, resulting in a malignant mechano-response. These findings reveal a previously under-appreciated mechanics-oncogene axis in colorectal cancer, in which physiological peristaltic forces cooperate with altered intracellular viscoelasticity to promote malignant cell states. This insight has important implications for CRC diagnosis, and treatment.

## Results

### Peristalsis-induced enrichment of LGR5^+^ cell populations in *KRAS* G12C mutated CRC cells and enhanced ERK signaling

We first evaluated if external application of peristalsis mechanics enhanced LGR5 stem cell populations within untransformed intestinal epithelial cells (HIEC-6) and KRAS G12C colorectal cancer (CRC) cells (SW837). LGR5 enrichment is an established read-out of peristalsis-associated malignant mechanoresponse^10,11^.

No significant changes were observed in LGR5 populations in untransformed intestinal epithelial cells, even with peristalsis stimulation (**Figure 1A**). However, in the *KRAS* G12C mutant cell line SW837, expectedly, peristalsis induced a significant ∼9X fold increase in LGR5 cancer stem cells compared to static controls (**Figure 1B**). Our data indicated that only cancerous cells with an oncogenic *KRAS* mutation transduced mechanical stimulation into a cancer stemness-promoting malignant mechano-response. Importantly, neither cell type demonstrated any increases in proliferation following peristalsis, evaluated via nuclear expression of the Ki67 antigen (**Supplementary Figure S1**).

**Figure 1.**
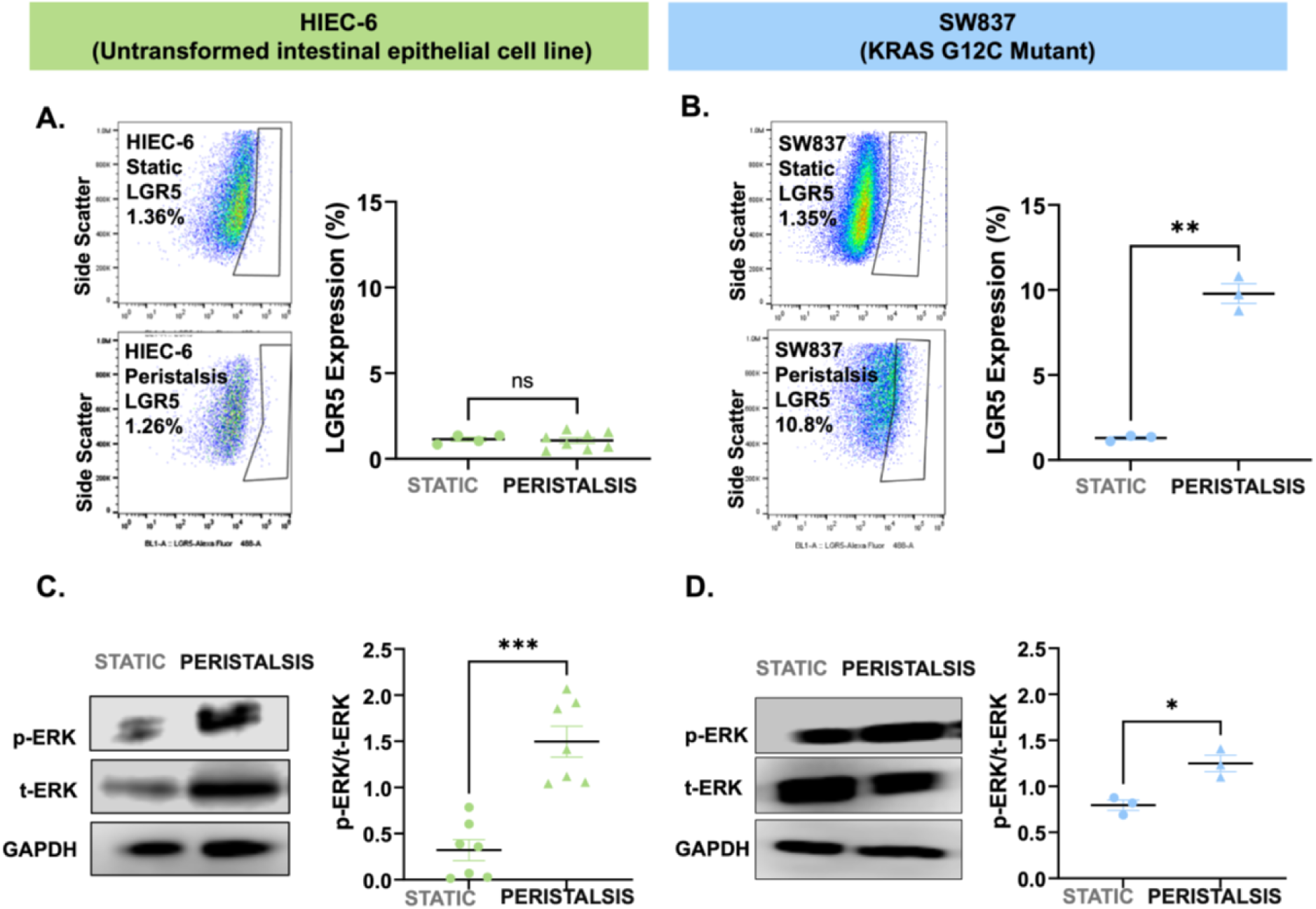
LGR5^+^ cancer stem cell enrichment and ERK activation in KRAS G12C mutant CRC cells. (A) Representative LGR5 flow cytometry plots of HIEC-6 cells maintained in static controls or exposed to peristalsis in the bioreactor for 24hrs. Individual dot plots summarizing flow analysis of HIEC-6 LGR5^+^ expression (%) after 24hr exposure to peristalsis bioreactor or maintenance in static controls (ns; not significant; t-test, n≥3). (B) Representative LGR5 flow cytometry plots of SW837 KRAS G12C cells maintained in static controls or exposed to peristalsis in the bioreactor. Individual dot plots summarizing flow analysis of KRAS mutant G12C LGR5^+^ expression (%) after 24hr exposure to peristalsis bioreactor or maintenance in static controls. Significant increase in LGR5^+^ expression was noted in G12C cells exposed to peristalsis compared to static controls. (**p<0.01, t-test, n≥3). (C) Representative western blots for phosphorylated ERK (pERK) activity in HIEC-6 cells after 24hr exposure to peristalsis or maintenance in static controls. Band intensities were normalized to their corresponding GAPDH loading control. Individual dot plots of densitometric quantification, reporting ratio of normalized p-ERK/t-ERK, demonstrating that peristalsis increased p-ERK levels compared to static controls in HIEC-6 cells. (***p<0.001, t-test, n≥3). The membranes were cut prior to antibody hybridization to process and visualize bands of interest. Full-length raw blot images, including all membrane edges and replicates, are provided in the supplementary information. (D) Representative western blots for phosphorylated ERK (pERK) activity in G12C cells after 24hr exposure to peristalsis or maintenance in static controls. Individual dot plots show densitometric quantification of p-ERK/t-ERK; peristalsis increased p-ERK levels compared to static controls in KRAS G12C mutated CRC cells. (*p<0.05, t-test, n≥3)

Next, we examined ERK pathway activation via phosphorylation (phospho-ERK), due to its central role as a KRAS effector, mechano-sensor, and maintenance of LGR5^+^ stem cells ^22,23^. Peristalsis increased ERK activation in untransformed HIEC-6 cells significantly (∼4.7X fold higher in peristalsis compared to static; **Figure 1C**). In *KRAS* G12C cells, baseline activity of phospho-ERK was already higher at static conditions due to constitutive KRAS oncogene activation (**Figure 1D**). Despite this elevation, peristalsis further amplified phospho-ERK activation additionally by 1.7X fold over static controls, consistent with KRAS pathway activation under mechanical stimulation.

Our findings demonstrated that both untransformed HIEC-6 cells and KRAS mutant SW837 cells were responsive to peristalsis mechanics by phosphorylating ERK. However, oncogenic KRAS was required to couple mechanical stimulation to an LGR5^+^ stemness promoting response, underscoring that *KRAS* mutations rewire how cells interpret mechanically activated signaling.

### *KRAS* G12C cells possess a distinct intrinsic mechanical phenotype

Given the divergent peristalsis associated mechano-response between untransformed HIEC-6 cells and SW837 KRAS mutant cells, we next examined if this was governed by differences in intrinsic cellular material properties using Brillouin microscopy (**Figure 2)**. Overlaid representative Brillouin heat maps are shown in **Figure 2A**, with visual differences between untransformed HIEC-6 cells and KRAS G12C mutant cells. Scattered photons were quantified for differences in Brillouin shift and Full Width Half Maximum (FWHM), corresponding to changes in intracellular elasticity and viscosity respectively (**Figure 2B**). Brillouin microscopy set up and how Brillouin Shift and FWHM readings are related to elasticity and viscosity is shown in **Supplementary Figure S2**^24^, with additional workflow examples in **Supplementary Figure S3**. Brillouin cellular measurements were averaged across the cell volume without distinguishing between intracellular structures. Comparisons between cells or conditions were therefore identified as a % change from control values. At baseline static conditions, in the absence of any external mechanical stimulation, KRAS mutant SW837 cells already had significantly elevated elasticity corresponding to a 2.71% increase in Brillouin shift vs. untransformed HIEC-6 cells. This was accompanied by 9% lower intracellular viscosity in KRAS mutant SW837 cells, compared to untransformed HIEC-6. Together, the higher elasticity and lower viscosity in KRAS mutant cells indicated a more agile and deformable cytoplasmic environment.

**Figure 2.**
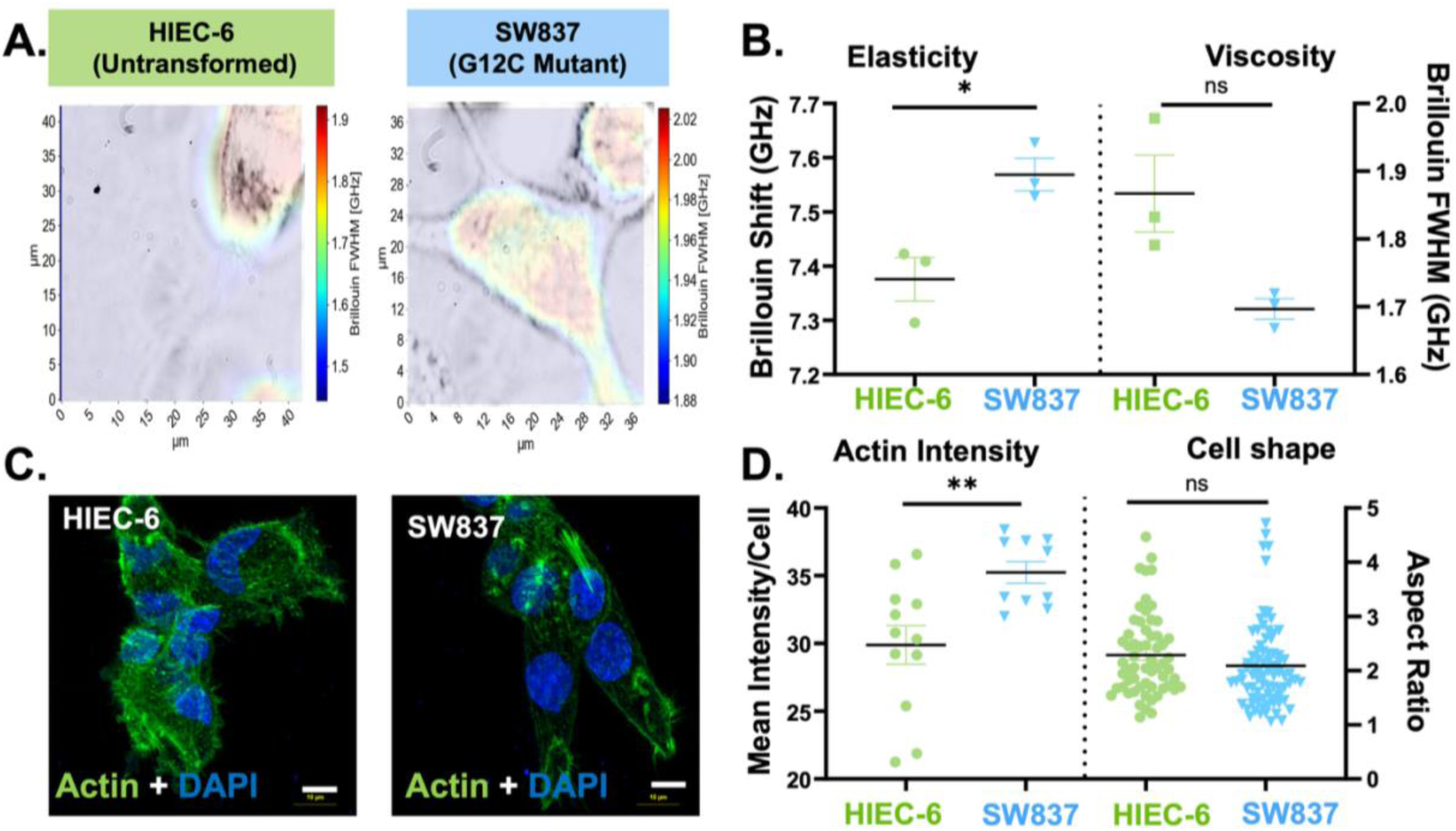
Cellular mechanical properties and morphologies differ between untransformed intestinal epithelial cells and *KRAS* G12C mutant colorectal cancer cells (A) Representative Brillouin images of HIEC-6 and SW837 KRAS G12C cells with an overlaid Brillouin shift and Full width half maximum (FWHM). (B) Quantification shows higher Brillouin shift and lower FWHM in SW837 vs. HIEC-6. (*p<0.05; ns = not significant; t-tests; n≥3). (C) Representative micrographs of HIEC-6 and SW837 cells stained with Phalloidin (green) and DAPI (blue). All scale bars 10 μM. (D) Individual dot plots quantifying mean actin intensity per cell and aspect ratio for both cell types. At baseline, KRAS G12C mutant SW837 had greater visualized actin compared to HIEC-6 cells. (**p<0.001; t-tests; n≥3), while cells were similarly elongated.

Visual differences in filamentous actin were also observed microscopically between HIEC-6 and SW837 cells (**Figure 2C**), quantified via actin intensity and cell shape differences between the two cell types (**Figure 2D).** At baseline, a 17.8% increase in visualized actin intensity via phalloidin staining was observed in SW837 cells compared to HIEC-6 cells (**Figure 2D**). This was not accompanied by visible or quantifiable differences in cell shape based on aspect ratios. Together, our data indicated that the increased deformability of KRAS mutant cells putatively governed the malignant mechano-response observed in **Figure 1**, although no major changes in visualized actin were observed.

### Peristalsis disrupts cellular mechanical homeostasis and cytoskeletal organization more readily in *KRAS* G12C compared to untransformed epithelial cells

Given the baseline differences in cellular deformability between KRAS mutant and untransformed epithelial cells, we next examined changes under external peristalsis stimulation. Cellular material properties were measured using Brillouin microspectroscopy under static or peristalsis stimulated conditions, showing representative Brillouin heatmaps and their associated quantifications for Brillouin shift and FWHM (**Figure 3**).

**Figure 3.**
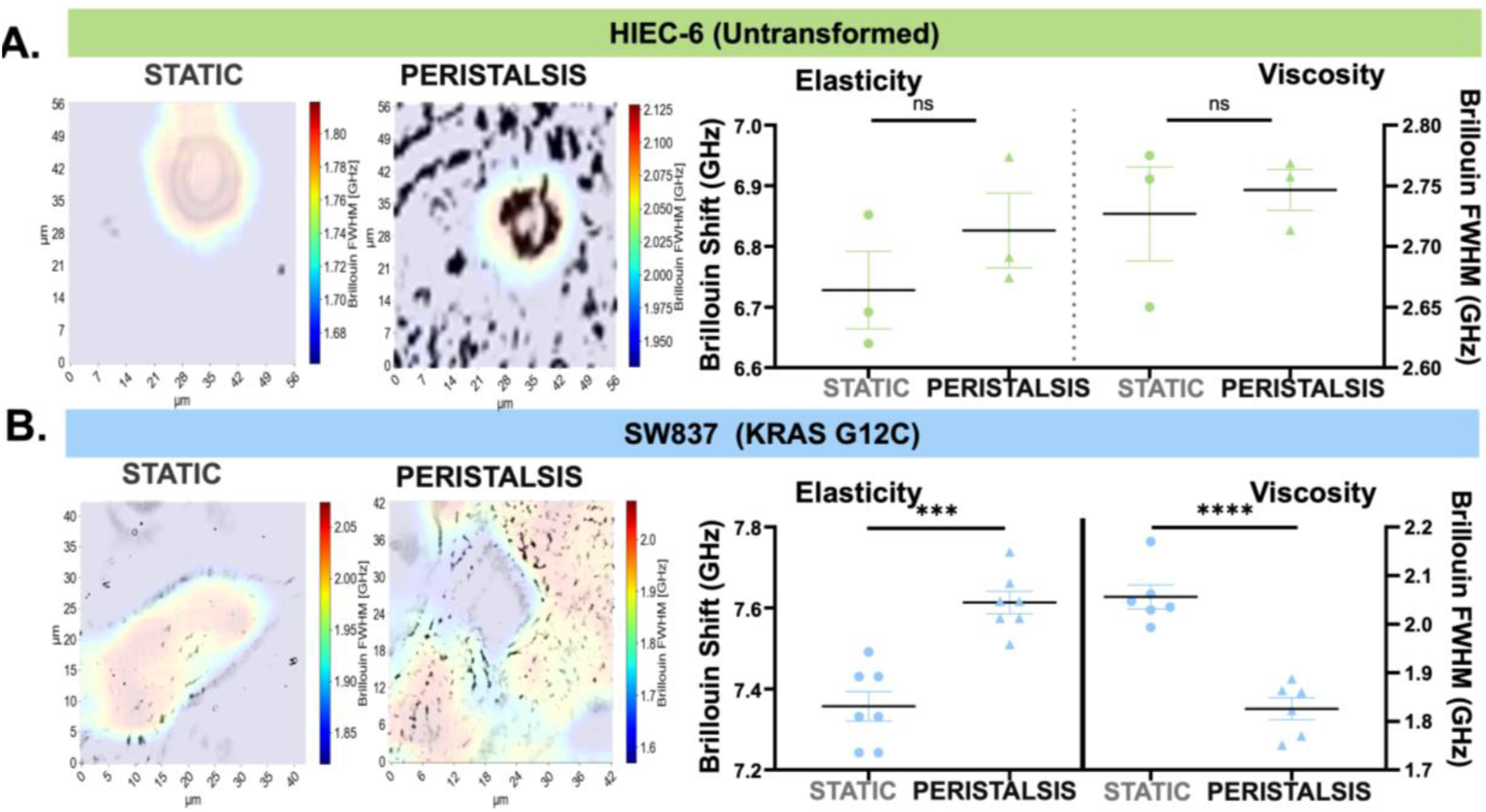
Peristalsis forces selectively amplify the viscoelastic properties of *KRAS* G12C cells. (A) Representative Brillouin images with an overlaid Brillouin shift and Full width half maximum (FWHM) in HIEC-6 cells under static controls or peristalsis exposure for 24hr; associated quantification of Brillouin shift and FWHM (ns; not significant; unpaired t-test, n=3). (B) Representative Brillouin images with an overlaid Brillouin shift and Full width half maximum (FWHM) in SW837 G12C under static controls or peristalsis exposure for 24hr. Quantification shows higher Brillouin shift and lower FWHM in SW837 G12C under peristalsis. (***p<0.001, ****p<0.0001; unpaired t-tests; n=3).

In response to peristalsis, untransformed HIEC-6 cells had unremarkable changes in elasticity and viscosity compared to static controls (**Figure 3A)**. In contrast, *KRAS* G12C cells exhibited a striking hyperresponsiveness to mechanical stimulation. Under peristalsis, Brillouin shift increased significantly by 3.47% over the static baseline (**Figure 3B),** further elevating elasticity. Simultaneously, peristalsis stimulation decreased FWHM starkly by 11.19% (**Figure 3B)**, indicating a marked reduction in intracellular viscosity. These changes in elasticity and viscosity following peristalsis stimulation amplified an already altered deformable state at baseline (**Figure 2)**, shifting KRAS mutant cells towards an even more deformable and instantaneous mechano-response. In contrast, untransformed non-cancerous epithelial cells maintained excellent mechanical homeostasis despite peristalsis stimulation.

As a consequence of the tight mechanical homeostasis in HIEC-6 cells, no significant changes in cytoskeletal organization, actin intensity or cell shape were observed between static and peristalsis-exposed cells (**Figure 4A**). Quantification of actin fiber alignment indicates minimal re-distribution in peristalsis-stimulated HIEC-6 cells, compared to their distribution in static controls (**Supplementary Figure S4**; HIEC-6). In contrast, peristalsis stimulation of the highly deformable of *KRAS* G12C cells resulted in pronounced cytoskeletal and morphological reorganization (**Figure 4B)**. Visually, actin organization was more apparent with peristaltic stimulation, quantified further via significant changes in alignment distrbutions (**Supplementary Figure S4;** SW837). This was additionally accompanied by a significant increase in visualized actin intensity per cell upon peristalsis (roughly 54% increase; *p<0.05, t-test; **Figure 4B)**. Visually, peristaltically stimulated KRAS G12C cells were also significantly more elongated compared to round cells maintained in static conditions. Cell elongation was reflected in the increased aspect ratio measure quantifying cell shape.

**Figure 4.**
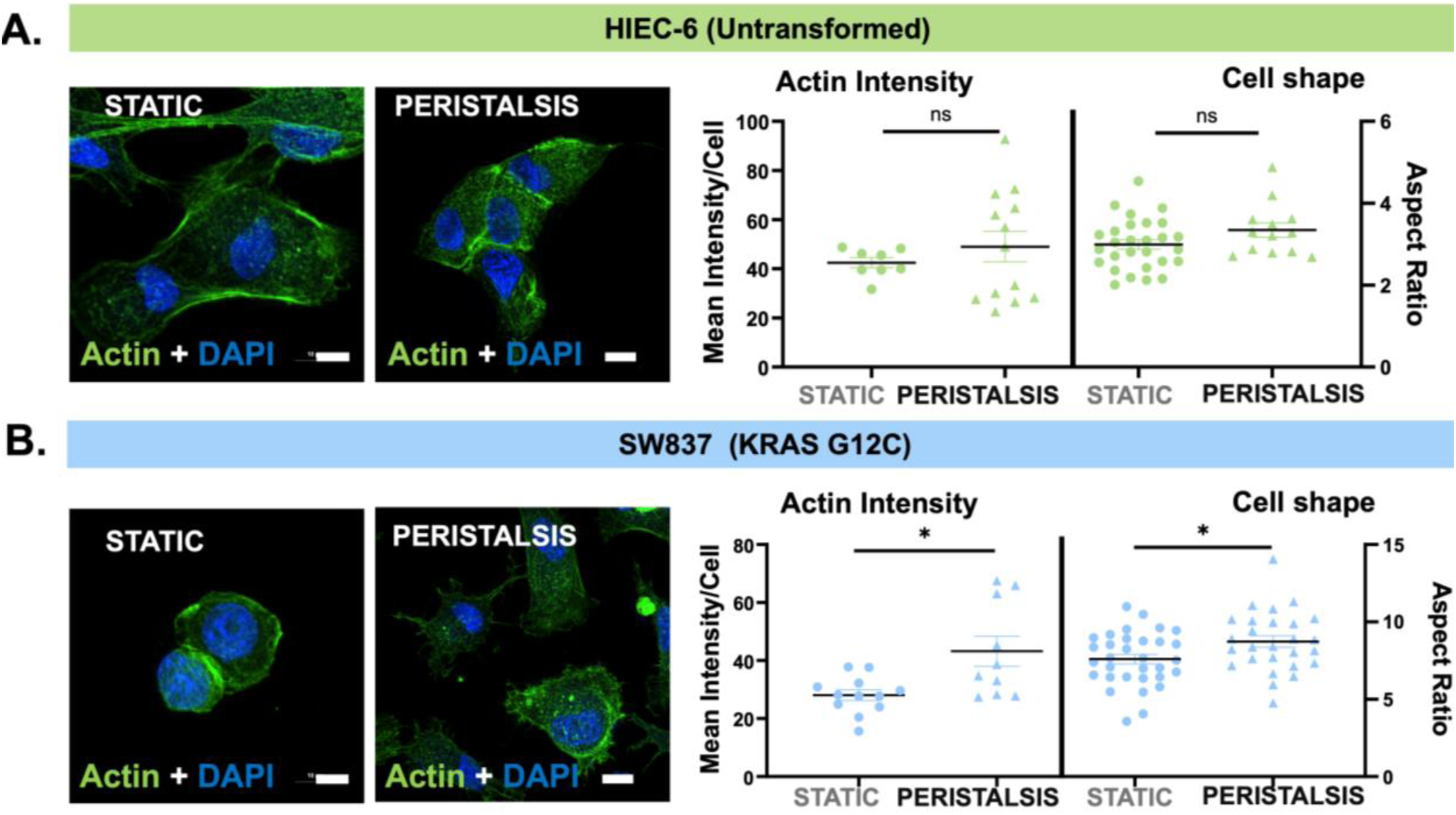
Peristalsis forces selectively increased actin intensity and elongated *KRAS* G12C cells (A) Representative micrographs of HIEC-6 cells stained with Phalloidin (green) and DAPI (blue) as static controls or following peristalsis for 24 hr. Individual dot plots quantifying mean actin intensity per cell and aspect ratio for HIEC-6 cells in static controls or following peristalsis (ns; not significant, t-test, n≥3). (B) Representative micrographs of SW837 *KRAS* G12C cells stained with phalloidin and DAPI as static controls or following peristalsis. Individual dot plots quantifying mean actin intensity per cell and aspect ratio for G12C cells. Actin intensity and aspect ratio increased significantly under peristalsis, compared to static controls (*p<0.05; t-tests; n≥3 Scale bar=10 μm).

### Inhibition of KRAS activity (RASi) suppressed peristalsis-driven mechanical remodeling and stemness

Given the hyper-responsiveness of KRAS G12C mutant cells to peristalsis, we next tested if KRAS activity was required to produce the outsized mechano-response using a covalent inhibitor, sotorasib (RASi). We first established a dose-response of KRAS G12C mutant cells in 2D cultures to identify a sotorasib dose that maintained viability of cells above 50%. This dose was established at 1.02 μM for KRAS G12C cells (**Supplementary Figure S5A).** This dose also effectively suppressed pERK activity by >50%, further underscoring its efficacy at successfully inhibiting KRAS effector activity (**Supplementary Figure S5B**). This concentration was used for KRAS activity inhibition for all subsequent experiments and are noted “RASi”.

Inhibition of KRAS activity completely mitigated peristalsis-induced deformability of SW837 cancer cells, implicating KRAS G12C activity as essential to driving the malignant mechano-response. In the presence of sotorasib inhibiting KRAS G12C activity (RASi), peristalsis failed to increase cellular elasticity. Instead, a 17.4% decrease in elasticity was observed (**Figure 5A)**. Likewise, peristalsis failed to reduce viscosity under RASi, non-significantly increasing it instead by 22%. These changes in cellular material properties with RASi implied reduced instantaneous deformability, which translated to the complete lack of peristalsis-induced cellular elongation **(Figure 5B)**.

**Figure 5.**
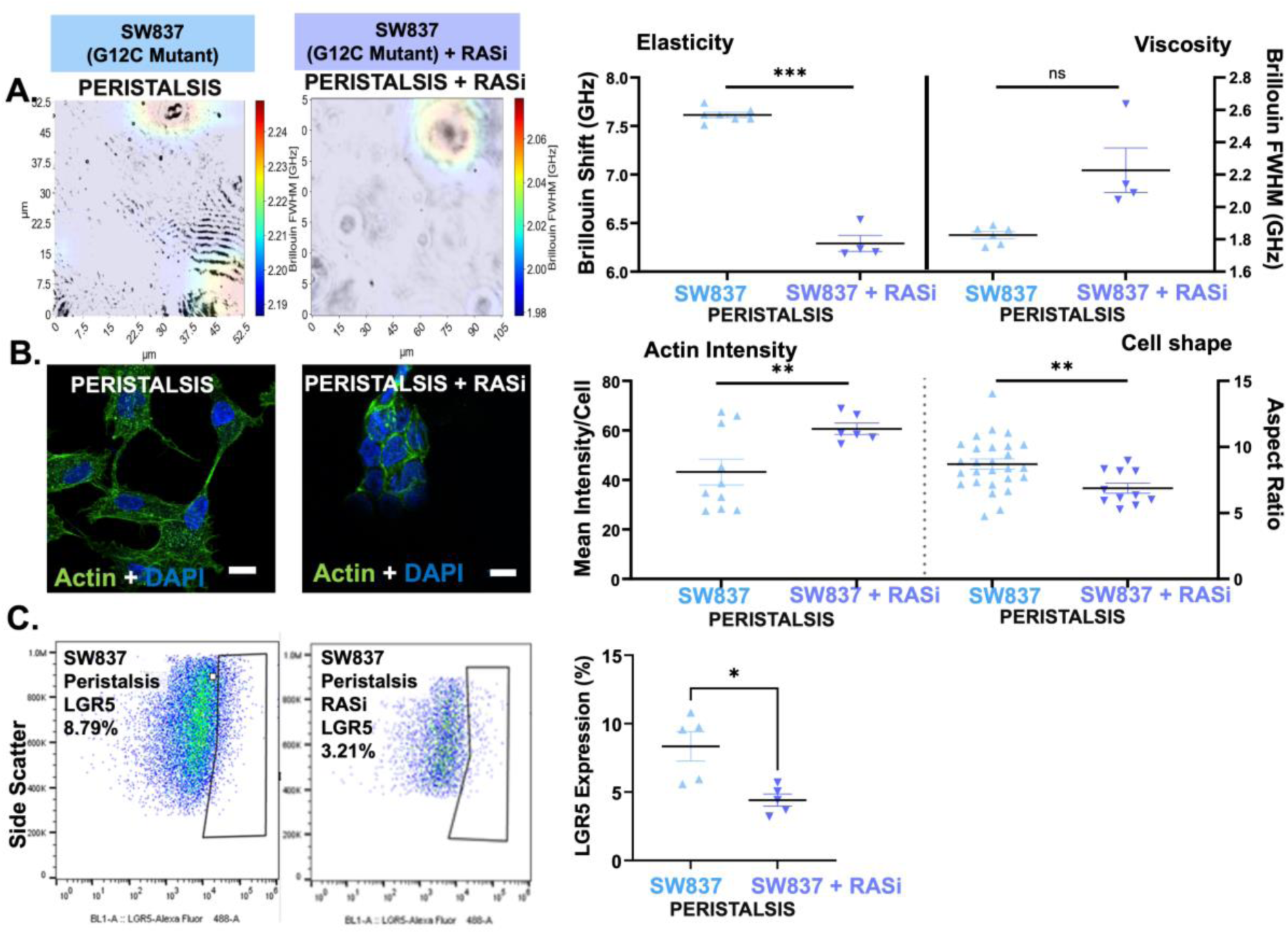
RAS inhibition (RASi) reverses *KRAS* G12C-driven material changes and peristalsis-associated malignant mechano-responses (A) Brillouin images with an overlaid Brillouin shift and Full width half maximum (FWHM) in SW837 G12C after peristalsis exposure for 24hr with/without RASi treatments; associated quantification via dot plots of Brillouin shift and FWHM in SW837 G12C mutant cells after peristalsis +/− RASi; (***p<0.001;ns – not significant; t-tests; n≥3). (B) Representative micrographs of SW837 G12C mutant cells stained with Phalloidin (green) and DAPI (blue) after peristalsis +/− RASi. Dot plots quantifying mean actin intensity per cell and aspect ratios following peristalsis +/− RASi. (**p<0.01; t-tests; n≥3) (C) Representative LGR5 flow cytometry plots of SW837 KRAS G12C cells exposed to peristalsis +/− RASi treatments, with associated quantification. Significant decrease in LGR5^+^ expression was noted in SW837 G12C cells in Peristalsis+RASi compared to uninhibited peristalsis. (*p<0.05, t-test; n≥3).

Lastly, the effect of RASi was tested on peristalsis-induced LGR5 enrichment. RASi alone did not decrease LGR5 in static controls in KRAS G12C SW837 mutant cells (**Supplementary Figure S6**). However, RASi expectedly curtailed peristalsis-induced LGR5 enrichment, which was lower by ∼50% compared to control peristalsis-induced conditions in the KRAS G12C mutant cells (**Figure 5C**). Together, our data strongly indicated that oncogenic KRAS activity was required to sustain the mechano-adaptive cell state and the associated malignant mechano-response induced by peristalsis.

### Ectopic expression of KRAS G12C in untransformed intestinal epithelial cells empowered a malignant mechanoresponse to peristalsis

Lastly, we investigated if a gain of KRAS G12C activity in an otherwise untransformed intestinal epithelial cell line (HIEC-6) may alter how they transduced peristalsis. HIEC-6 cells were transfected to transiently express the KRAS G12C oncogenic protein. Expression of KRAS G12C was validated using immunoblots for increased expression of RAS, demonstrating that transfection resulted in a 2-fold increase in RAS protein expression compared to HIEC-6 control cells (*p<0.05, t-test; **Supplementary Figure S7**).

Interestingly, introduction of KRAS G12C activity into HIEC-6 meant that it no longer maintained mechanical homeostasis in response to peristalsis stimulation. Following peristalsis, significant shifts in cellular elasticity and viscosity occurred in HIEC-6 G12C mutant cells (**Figure 6A)**. This shift in cellular deformability was also accompanied by significant increases in actin intensity (**Figure 6B**), and significant cell elongation quantified via increased aspect ratio (**Figure 6B**). Lastly, exposure to peristalsis in HIEC-6 G12C cells led to significant cancer stem cell LGR5 enrichment (**Figure 6C**), over untransformed peristaltically stimulated HIEC-6 cells. No differences were observed in levels of LGR5 between statically maintained HIEC-6 G12C cells, compared to wild type HIEC-6 cells (**Supplementary Figure S8**), implying that peristalsis stimulation and oncogenic transformation were both necessary to induce LGR5 enrichment. These results further underscored that the expression of the KRAS G12C oncoprotein in non-cancerous intestinal epithelial cells sufficiently phenocopied the malignant mechano-responses observed in KRAS G12C mutant colorectal cancer cells.

**Figure 6.**
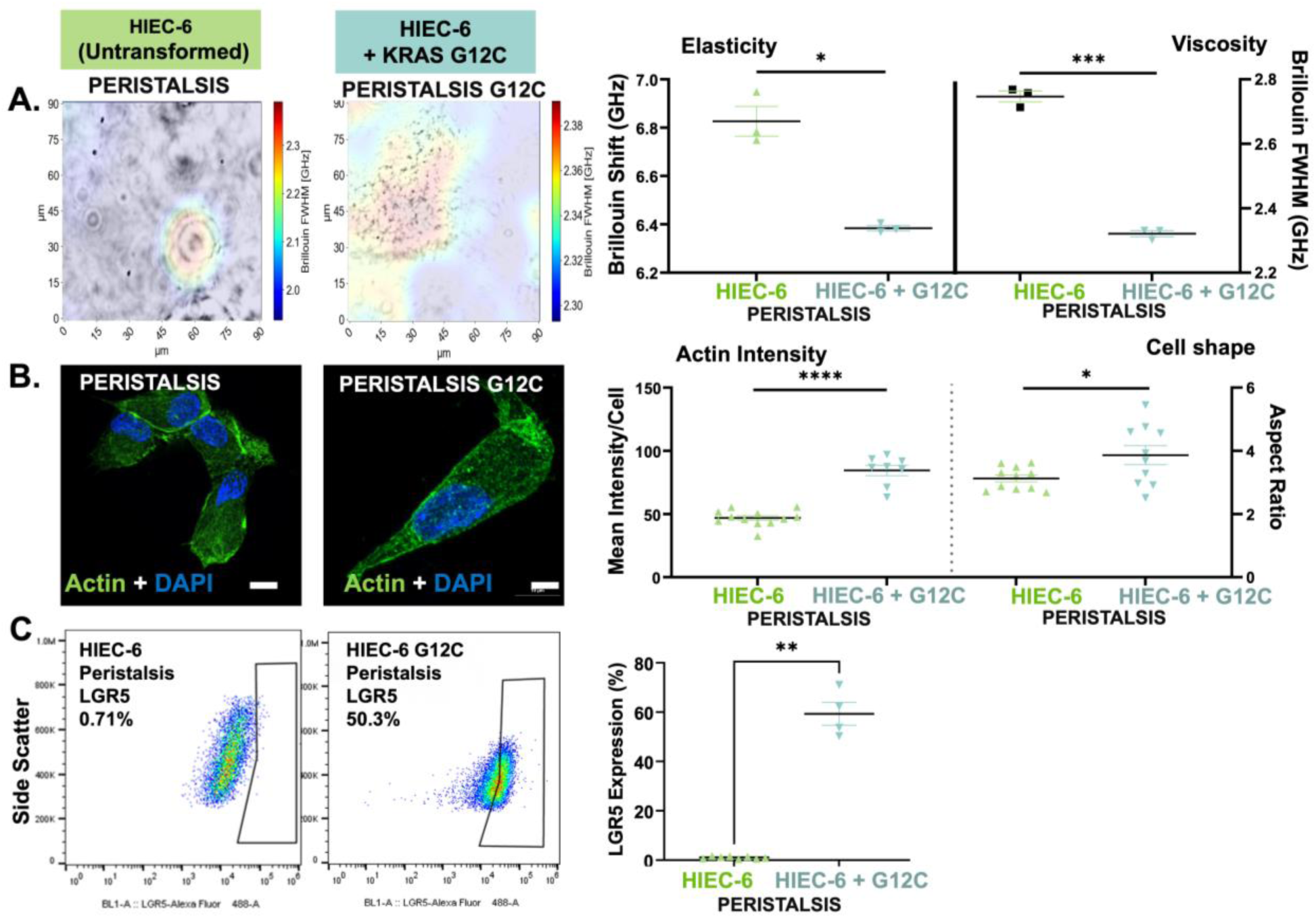
KRAS G12C transformed HIEC-6 cells led to peristalsis-induced malignant mechano-responses. (A) Representative Brillouin images with an overlaid Brillouin shift and FWHM in HIEC-6 +/− KRAS G12C mutation following peristalsis, with associated quantification. (*p<0.05, ***p<0.001; t-test; n≥3). (B) Representative micrographs of HIEC-6 cells +/− KRAS G12C mutation stained with Phalloidin (green) and DAPI (blue) after peristalsis exposure, with quantification of actin intensity per cell and aspect ratio; (****p<0.0001, *p<0.05, t-test; n≥3) (C) Representative LGR5 flow cytometry plots of HIEC-6 cells +/− KRAS G12C mutation following peristalsis, with associated quantification of LGR5 expression. Significant increase in LGR5^+^ expression was noted in HIEC-6 G12C cells exposed to peristalsis compared to peristalsis mechanical stimulation. (**p<0.01, t-test; n≥3).

## Discussion

Colorectal cancer develops within a uniquely dynamic mechanical environment, as the colonic epithelium is continuously exposed to rhythmic deformation from colonic peristalsis ^25^. Our work strongly demonstrated that a malignant mechano-response downstream of peristalsis was confined to cells with activated oncogenic KRAS. In this current work, we address the fundamental ‘why’ underlying this divergent malignant mechano-response by examining cellular material and mechanical properties. We focused on mechanistically dissecting the *KRAS* G12C mutant variant, due to its clinical actionability via a covalent inhibitor, sotorasib^26^. Although *KRAS* G12C occurs in only ∼3% of CRC patients, it is associated with aggressive tumor behavior and reduced survival^27,28^. Comparing mechano-responses across untransformed intestinal epithelial cells and *KRAS* G12C cells, our results re-iterated that peristalsis-associated malignant mechano-response was confined to cells with an active *KRAS* oncogene (LGR5 cancer stem cell increases in **Figure 1**). Although the present study focuses on *KRAS* G12C, previous work has demonstrated peristalsis-induced LGR5 enrichment in other *KRAS* mutant variants in colorectal cancer^10^, suggesting that mechano-sensitivity may represent a broader feature of oncogenic KRAS signaling rather than a G12C-specific phenomenon. The increased phosphorylation of ERK downstream of mechanics demonstrated that both untransformed healthy cells and cancer cells sensed and responded to peristalsis mechanics. Despite this, the divergent mechano-response suggested that oncogenic KRAS rewired how epithelial cells interpreted mechanical cues, potentially by altering intrinsic cellular material properties that govern mechanical responsiveness.

We used Brillouin microscopy to quantify intracellular viscoelasticity. Unlike traditional mechanical assays such as atomic force microscopy or nanoindentation, which require direct indentation or manipulation^29^, Brillouin microscopy enables non-contact, non-destructive assessment of intracellular mechanical properties. In doing so, Brillouin imaging provides insights on cellular elasticity (Brillouin shift) and cellular viscosity (Brillouin full width half maximum; FWHM)^30^. Using Brillouin, our work identified that baseline mechanical properties of SW837 *KRAS* G12C cancer cells were starkly different (more elastic, less viscous) compared to untransformed HIEC-6 intestinal epithelial cells (**Figure 2**). These results support the notion that tumor cells exhibit altered mechanical phenotypes relative to normal counterparts^15,31,32^. Importantly, these baseline differences were accompanied by elevated actin intensity in KRAS G12C cells, consistent with literature showing that KRAS-driven cytoskeletal remodeling and actin reorganization contribute to mechanical changes and deformability in cancer cells^33,34^.

Upon exposure to peristalsis, *KRAS* G12C cells exhibited an exaggerated mechanical response, with further increased elasticity and decreased viscosity (**Figure 3)**, indicating a shift toward a hyper-deformable, mechanically responsive state. In contrast, HIEC-6 cells showed minimal changes and largely maintained mechanical homeostasis. This difference supports the idea that oncogenic transformation compromises mechanical stability, allowing external forces to induce cytoskeletal changes rather than controlled homeostatic compensation. This is also consistent with reports that oncogenic *KRAS* activation changes the dynamics of actin remodeling and intracellular tension through its many effector pathways and other mechanosensitive pathways like ERK and YAP/TAZ^35^. Consistent with this interpretation, peristalsis induced striking actin reorganization in *KRAS* G12C cells, including aligned parallel fiber structures, along with increased cell elongation (**Figure 4B)**. Cell elongation and actin reinforcement are both correlated to migratory and invasive behavior^36^, suggesting that peristalsis not only alters intracellular material properties but also promotes morphologic phenotypes linked to invasiveness.

A limitation of the baseline and peristalsis-induced comparisons is that SW837 and HIEC-6 cells are non-isogenic models that differ in numerous genetic and phenotypic characteristics beyond just KRAS status. Therefore, the differences observed under baseline and peristaltic conditions cannot be attributed solely to KRAS G12C. To address causality, we employed both pharmacologic inhibition of KRAS activity and ectopic over-expression of KRAS G12C to investigate if KRAS activity was necessary for the observed mechanosensitive responses.

Our findings establish the *KRAS* G12C oncogene as a primary driver of cellular material properties, fundamentally altering how cells respond to mechanical stimuli. This role was confirmed through two distinct approaches: (1) inhibition of KRAS G12C activity, which reversed the agile and deformable cellular mechanotype (**Figure 5**); and (2) the expression of the oncogene in untransformed epithelial cells, which decreased cellular viscosity and induced significant deformation and LGR5 enrichment (**Figure 6**). Despite employing transient transfections in the gain of function studies on untransformed HIEC-6 cells, our data strongly demonstrated increased RAS protein expression at the 72hr time point, where subsequent downstream analyses of mechano-responses were performed. Future studies may consider inducing more stable, population-selectable methods of RAS oncoprotein expression to enable longer-term mechanistic studies. Importantly, neither KRAS inhibition under static conditions in SW837 G12C cells nor KRAS G12C overexpression in untransformed HIEC-6 cells was sufficient to increase LGR5 levels at baseline in the absence of mechanical stimulation. This strongly indicated that oncogenic KRAS requires an external stimulus in the form of mechanical cues to drive LGR5 enrichment.

A central aspect of this KRAS-driven mechanotype is the reduction of viscosity associated with higher oncogenic activity. To the best of our knowledge, this phenomenon has not been previously reported in the context of colorectal cancer mechanobiology, especially in response to external mechanical forces. While recent studies have linked decreased membrane viscosity to migrating colorectal cancer cells ^37^, or changes in cytoplasmic viscosity to macromolecular crowding ^38^, these findings were not established in the context of a mechanoresponse, or specific to intracellular viscosity.

Ultimately, our results implicate a KRAS oncogene-mechanics axis that transforms the cellular mechano-type in response to peristalsis, likely over-riding the homeostatic visco-adaptive mechanisms typically found in healthy epithelial cells ^39^. While the exact pathways by which oncogenic KRAS alters cellular viscosity remain uncharted, existing evidence provides some candidates for consideration. For example, increased mTOR complex activity, a direct result of KRAS signaling and peristalsis, is known to alter cellular viscosity ^40^. Additionally, mechano-sensitive aquaporin channels may facilitate water influx that dilutes the cytoplasm, thereby reducing viscosity in response to physical forces of peristalsis ^41,42^.

Despite these interesting mechanistic angles, these insights should be interpreted considering several model-related limitations including the limited number of cell lines tested. Furthermore, HIEC-6 cells are derived from fetal intestinal epithelial cells and do not fully recapitulate the mature colonic epithelium, despite being a widely acceptable model to study the intestinal epithelium. Future validation will prioritize additional oncogenic KRAS G12C lines including patient-derived models, and isogenic models to establish the translational relevance of these findings.

## Conclusion

In summary, our study defines a novel KRAS oncogene-mechanics axis that alters the biophysical state of cells. By lowering intracellular viscosity, oncogenic KRAS G12C creates a hyper-deformable mechano-phenotype, allowing cells to leverage, rather than resist, the mechanical forces of peristalsis. Our findings demonstrate that KRAS driven colorectal cancer synergizes with the mechanically dynamic colon microenvironment by altering cellular material properties. This opens out new ways to target the mechanical state or mechanotransduction to complement emerging KRAS-directed therapies.

## Acknowledgements and Funding Statement

This work was supported by grants from the Cancer Prevention and Research Institute of Texas RP230204 (SAR) through the Regional Excellence Center in Cancer program at Texas A&M University. This work was additionally supported by NCI R37CA269224 (SAR). The authors acknowledge support from Dr. Taylor H. Ware and Dr. Manivannan Sivaperuman Kalairaj for assistance with bacterial cultures to amplify the plasmid used in this study.

## Data Availability Statement

Raw data associated with this study are deposited on the Raghavan Lab Dataverse in the Texas Data Repository with the doi: https://doi.org/10.18738/T8/W2Q0YN; additional supporting data is also presented in supplementary information.

## Author Contributions

A.L. Formal analysis, Investigation, Methodology, Software, Validation, Visualization, Writing-original draft, review and editing.

V.C. Formal analysis, Investigation, Methodology

M.K. Formal analysis, Investigation, Methodology, Visualization

A.S. Methodology, Visualization

A.H. Methodology, Visualization

V.Y. Conceptualization, Resources, Investigation, Project Administration, Supervision, Writing – review and editing

S.A.R. Conceptualization, Funding Acquisition, Resources, Investigation, Visualization, Project Administration, Supervision, Validation, Writing – original draft and review and editing.

All authors reviewed and approved the manuscript.

## Materials and Methods

### Materials

Cell culture reagents were purchased from ThermoFisher Scientific (Waltham, MA) unless otherwise specified. All cell lines were purchased from American Type Culture Collection (ATCC; Manassas, VA) unless otherwise specified. Polydimethylsiloxane (PDMS) was purchased from DOW Chemical (Midland, MI). All other chemical reagents were purchased from Sigma Aldrich (St. Louis, MO), unless otherwise indicated. Antibodies used for cellular staining were purchased from Santa Cruz Biotechnology (Dallas, TX) unless otherwise indicated.

### Cell Culture

Colorectal cancer cell line SW837 and untransformed intestinal epithelial cell line HIEC-6 were used in this work. SW837 was used before passage 20 and HIEC-6 cells were used before passage 8. Dulbecco’s Modified Eagle Medium (DMEM) supplemented with 10% heat-inactivated fetal bovine serum (Peak Serum, Inc., Wellington, CO) and 1X Antibiotic-Antimycotic solution was used as the primary growth media for SW837 cells. Opti-Minimal Essential Medium (Opti-MEM) reduced serum medium supplemented with 4% heat-inactivated fetal bovine serum (Peak Serum, Inc., Wellington, CO), 20 mM HEPES, 10 mM Glutamax and 10 ng/ml Epidermal Growth Factor (EGF) was used as the primary growth media for HIEC6 cells. All cells were cultured in standard 2D tissue culture flasks and treated with 0.25% trypsin to dissociate adherent cells. All cultures were routinely screened to ensure cultures were free of mycoplasma.

### Preparation of Cell-Seeded Membranes and Bioreactor Assembly

Polydimethyl siloxane (PDMS) membranes were prepared at a 10:1 ratio using previously established protocols^9^. To enhance cell attachment to PDMS, the membrane seeding areas were coated with Collagen I (200 μg/ml). Cells were plated onto the PDMS at a monolayer density of 250,000 cells/mL and incubated at 37°C for 24 hours to allow adhesion. Cell attachment was confirmed by visual inspection using a cell culture microscope.

Operation of the peristalsis bioreactor followed previously established methods^9^. Briefly, cell culture medium was removed from each cell-seeded PDMS membrane after 24 hours. Cell-seeded peristalsis membranes were then placed into the peristalsis bioreactor bottom while static membranes were maintained in cell culture dishes with 1 mL of fresh media. The bioreactor was secured with commercially available zip ties and connected to the peristalsis pump. The assembled set up was transferred to a 37°C incubator and connected to an Arduino preprogrammed code to apply 0.4 Pa shear stress and 15% cyclic strain at 12 rpm^9^, mimicking the range of forces found in colonic peristalsis^43,44^. Cells were incubated at 37°C for 24 hours, either maintained as static controls or subjected to peristaltic stimulation in the bioreactor and were then collected for downstream analysis.

Control (untreated) samples were maintained in static membranes, or within peristalsis bioreactors in their respective growth medium. Media was supplemented with a sub-IC50 dose of the RAS inhibitor (sotorasib; 1 μM for SW837 identified in **Supplementary Figure S5)** to evaluate the effect of RAS inhibition on peristalsis-associated mechanotransduction.

### Flow Cytometry Assessment of LGR5^+^ Cancer Stem Cells

After 24 hours under either static or peristaltic conditions, cells were detached with 0.25% trypsin and recovered from the PDMS as single-cell suspensions in FACS buffer (Phosphate Buffered Saline (PBS) supplemented with 2% fetal bovine serum). Flow cytometry was carried out following previously optimized protocols for cancer cells^45,46^. Cells were incubated for 30 minutes at 37°C with AlexaFluor488-conjugated anti-LGR5 antibody (R&D Systems) or an isotype-matched AlexaFluor-488 control (R&D Systems), then washed, resuspended in fresh FACS buffer, and analyzed on an Attune NxT flow cytometer (ThermoFisher Scientific). Isotype controls defined a background gate at 0.5%, and the proportion of LGR5-positive cells was calculated and compared between static and peristalsis conditions. Gating strategy and isotype controls for each plot is demonstrated in **Supplementary Figure S9**. Identical gate coordinates were applied within a given cell type and experimental conditions. Gates were established using each experimental run’s corresponding controls and maintained throughout analysis to minimize operator bias. Isotype gates were cut off under ∼0.5% and applied as is, to determine differences in LGR5 levels.

### Quantification of Actin Intensity and Cell Shape

Cell-seeded PDMS membranes for SW837 and HIEC-6 cells from static, mechanically stimulated, and molecularly stimulated conditions were trimmed along the cell seeding area, and fixed in 4% formalin for 15 minutes at room temperature. Samples were permeabilized and blocked at room temperature for 1 hour in a solution of 0.15% Triton X-100 with 5% fetal bovine serum (FBS). Following blocking, cells were incubated for 1 hour at room temperature with fluorescently tagged AlexaFluour488-Phalloidin to visualize actin filaments (green). A nuclear counterstain, DAPI (blue), was included during this antibody incubation. Unbound probes were removed by rinsing with PBS, and the cell-seeded PDMS membranes were mounted using an antifade mounting medium. Fluorescent imaging was performed on a Nikon AX R point-scanning confocal microscope system (Nikon Instruments Inc., NY, USA), with five independents, non-overlapping regions imaged per sample for analysis. Fluorometric quantification was carried out using NIH ImageJ.

Mean actin intensity and cell shape measures (aspect ratio) were quantified from calibrated, fluorescently visualized actin using ImageJ (analysis workflow visualized in **Supplementary Figure S10).** Laser power, gain, and exposure time were held constant across all conditions. For each cell, cell elongation was determined by tracing the cell’s major axis to obtain a length value in micrometers (µm). Several cells were measured across five non-overlapping images, from at least three independent experimental runs.

### Brillouin Spectroscopy

Brillouin spectroscopy was used to assess the material and mechanical properties of cells to assess viscoelastic properties of the cell with/without mechanical stimulation^47^. To determine the effect of peristalsis on the mechanical properties of the cells, cells that had been maintained under static conditions or subjected to peristaltic stimulation for 24 hours were used. Cells seeded on the PDMS membrane were trimmed along the cell seeding area and put on a glass bottom slide. The slide was securely positioned on the microscope stage and remained fixed throughout all experimental conditions conducted during the day.

All samples were imaged within the glass bottom slide, where the cells were adhered to the PDMS membrane and submerged in the nutrient media. Viscoelasticity data were acquired using a custom-built upright confocal Brillouin microscope configured in a backscattering geometry^30^. A detailed description of the instrument and its operation have been reported previously^48,49^. A schematic of the spectrometer is shown in **Supplementary Figure S2** followed by a description of its operating principles. Brillouin cellular measurements were averaged across the entire cell volume without distinguishing between intracellular structures such as the nucleus and cytoplasm using methods established previously^30^. Briefly, cell segmentation was performed using an automated algorithm to identify individual cells within each image frame, and any subsequent analysis was restricted to pixels within the segmented cell boundaries. An example workflow illustrating Brillouin microscopy analysis providing whole cell viscoelastic properties is shown in **Supplementary Figure S3.** This approach yields a whole-cell average of the Brillouin shift and Brillouin line width, quantified via the full width half maximum (FWHM). To compare viscoelasticity between experimental conditions, average Brillouin Shift or FWHM were compared as changes between each condition, and represented as a % change, where indicated.

### Western Blot Analysis

Cells maintained under static conditions or subjected to peristaltic stimulation for 24 hours were detached with 0.25% trypsin, recovered from the PDMS, and lysed in 100 µL of radio-immunoprecipitation assay (RIPA) buffer supplemented with 1 µL of Halt Protease Inhibitor Cocktail. Total protein in the extracts was quantified using the Pierce™ BCA Protein Assay Reagent according to the manufacturer’s instructions for a 96-well format. After protein determination, 10 µg of protein from each sample were loaded onto 4–20% gradient polyacrylamide gels (Novex™, ThermoFisher) and separated via electrophoresis. Resolved proteins were transferred to a PVDF membrane, which was blocked with 2.5% bovine serum albumin (BSA). Primary antibodies were phospho-p44/42 MAPK (pErk1/2, Cat. No 9101), p44/42 MAPK (Erk1/2, Cat. No 9102), and GAPDH (Cat. No 5174) (Cell Signaling Technology). Solutions of primary antibodies were prepared at concentrations recommended by the manufacturer and used. Membranes were probed with primary antibodies overnight at 4°C, washed with TBST buffer, and probed with an appropriate HRP-conjugated secondary antibody for 1h, followed by repeated washing. GAPDH was used as a loading control. Detection was carried out using ECL chemiluminescence detection kit (SuperSignal TM West Pico PLUS Chemiluminsecent Substrate, Thermo Scientific) and images using LI-COR® 3600 C-Digit Blot Scanner. Densitometry was performed using NIH Image J. Band intensities were normalized by dividing the intensity obtained from each protein band to their corresponding loading control band intensity. Uncropped western blots are shown in **Supplementary Figure S11**. The original membrane was sectioned prior to antibody hybridization. All available raw blot images, including the uncropped regions for the main figure panels, are included in the Supplementary information. All blots were processed in parallel, and only representative blots are shown in the main figures. Raw blots for each representative blot are included in the supplementary which includes visible ladder bands and the membrane edges.

### KRAS G12C Expression in HIEC-6 Cells

pcDNA *KRAS* G12C plasmid was purchased from Addgene (Cat. No. 224284) and amplified in EColi. Plasmid DNA was isolated from the resulting culture using a standard miniprep procedure, and DNA concentration and purity were assessed spectrophotometrically prior to transfection. HIEC6 cells were maintained under standard culture conditions and seeded in 24-well plates to achieve optimal confluency for transfection. HIEC-6 cells were transiently transfected with *KRAS* G12C plasmid DNA using Lipofectamine 2000 (Thermo Fisher Scientific) according to manufacturer’s instructions for 24-well formats. Transfected cells were maintained under standard culture conditions until downstream analyses at 72hrs. To validate expression of the *KRAS* G12C construct, a sample of cells was lysed in RIPA buffer, for subsequent western blot analysis of RAS protein expression (Catalog No. 91054, Cell Signaling Technologies), with untransformed intestinal HIEC-6 cells serving as controls.

### Statistical Analysis

Statistical analysis was performed on GraphPad Prism 10. All reported values are means ± SEM, n and result from at least 3-5 independent biological replicates. Image analysis and morphometry included 3-5 non-overlapping fields of view from 3-5 biological replicates. Hypothesis testing was performed using GraphPad Prism, with Welch’s corrections applied to unpaired t-tests where appropriate. Statistical significance is indicated within each experiment with associated p-values. Individual dot plots were used to represent collected data, with a line at mean and error bars across the standard error of the mean.

## Supplementary Information

Supplementary files include unprocessed data or data processing steps that support the conclusions in the main text; these include Brillouin set up and analyses workflow, flow cytometry gating strategy, uncropped western blots and workflows determining dose of RAS inhibition and actin-based aspect ratio and alignment.

