## Supplementary Information for "Oncogene-Mechanics Axis: KRAS G12C Confers Agility Enabling Malignant Mechano-responses to Peristalsis in Colorectal Cancer"

**Supplementary Material**

**S1. Ki67 expression in HIEC-6 and SW837**

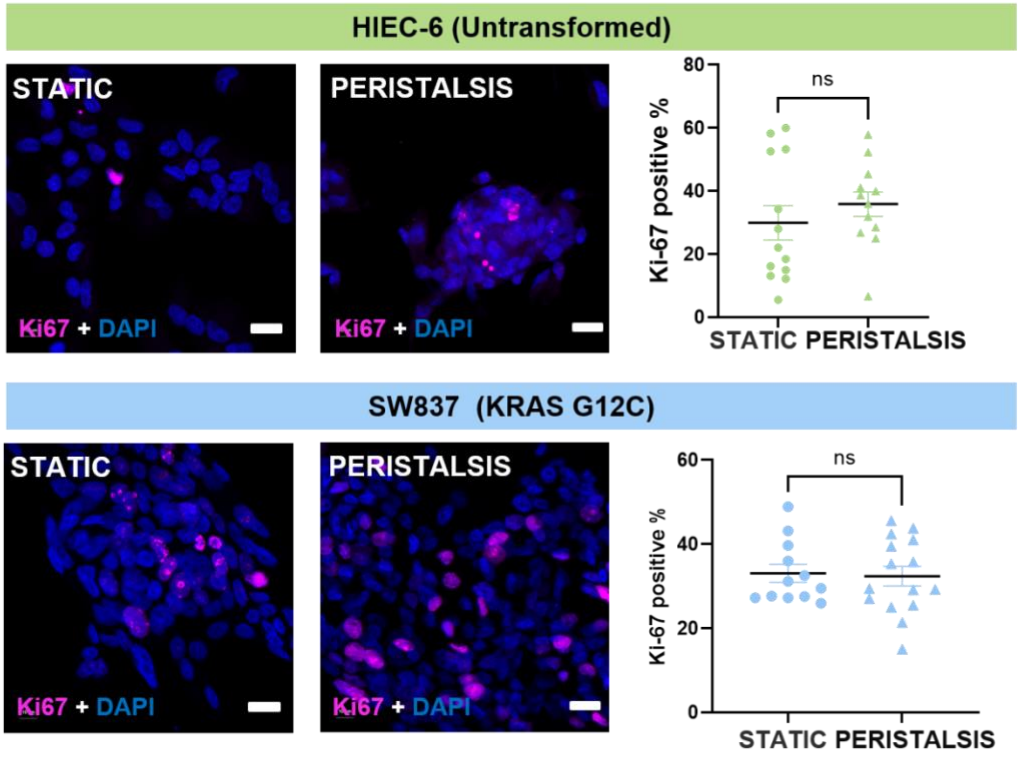


**Figure S1.** Ki67 immunofluorescent staining of (A) HIEC-6 untransformed cell (B) SW837 KRAS G12C exposed to peristalsis. Representative fluorescence micrographs stained with Ki67 antigen (magenta fluorescence) and counterstained with nuclear marker, DAPI (blue fluorescence). Scale bar 10 µm. Cells were maintained as static controls or exposed to 24h of peristaltic forces. Individual dot plot showing KI-67–positive cell percentages under static and peristaltic conditions. Each dot represents one sample; horizontal bars indicate mean ± SEM. ns, t-test, n≥3.

**S.2 Brillouin microscopy workflow**

A single-frequency laser was used to probe Brillouin-scattered photons. The excitation beam was directed to the sample using a set of polarization-control optics. By adjusting the half-wave plate (HWP), the optical power delivered to the sample through the microscope objective (MO) was optimized for imaging. During all imaging experiments, the incident power at the sample did not exceed 5 mW to avoid sample degradation.

Brillouin-scattered photons were collected by the same MO used for excitation and directed toward the confocal detection path. A quarter-wave plate (QWP) was used to rotate the polarization, enabling efficient routing of the collected signal toward the detection optics.

Optical sectioning and improved axial and lateral resolution were achieved using a confocal pinhole assembly (L2, PH, L3). The spatially filtered signal was then directed to a single-stage virtually imaged phased array (VIPA) spectrometer equipped with a hyperfine spectral filter (VC). The signal was focused on the VIPA etalon, where it underwent multiple-beam interference. The resulting interference pattern was Fourier transformed by a plano-convex lens (L4) and imaged onto the electron-multiplying CCD (EMCCD) detector.

Alignment and verification of the laser waist position were performed using an integrated wide-field microscope module consisting of an LED illumination source and a CMOS camera. The operating mode of the system (Brillouin spectroscopy or wide-field imaging) was selected by switching a broadband beam splitter (BBS).

The EMCCD records four replicated spectra, which are used to correct for the nonlinear spectral dispersion of the VIPA spectrometer and convert the measured peak positions into Brillouin frequency shifts.

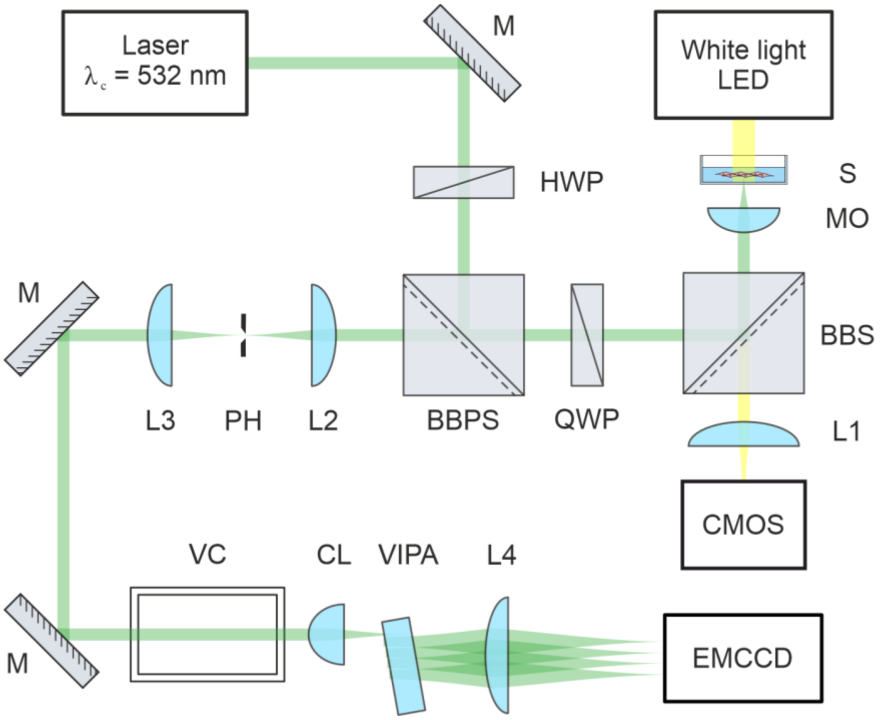


**Figure S2**. Confocal Brillouin spectrometer schematic. Here M – dielectric mirror, HWP – half-wave plate, BBPS – broadband polarizing beam-splitter cube, QWP – quarter-wave plate, BBS – broadband beamsplitter cube, MO – microscope objective, S – sample, L – plano-convex lens, CMOS – camera with complimentary metal-oxide-semiconductor sensor, PH – precision pinhole aperture, VC – molecular Iodine vapor cell, CL – cylindrical lens, VIPA – virtually imaged phased array, EMCCD – CCD camera with electron multiplication capabilities.

Brillouin microscopy relies on inelastic scattering of light from thermally driven density fluctuations, providing direct access to high frequency viscoelastic response. Analysis of the scattered spectrum yields a frequency shift, $\Omega$, and linewidth (at the full width half maximum, or FWHM), $\Delta$, which encode the real and imaginary components of the complex longitudinal modulus. Specifically, the storage (elastic) modulus is given by $M' = \rho(\lambda\Omega/2)^2$, where $\rho$ is the material density, $\lambda$ is the optical wavelength in the medium, while the loss (viscous) modulus follows as $M'' = M'(\Delta/\Omega)$. These quantities probe mechanical behavior in the GHz regime.

#### S.3 Brillouin microscopy analysis workflow

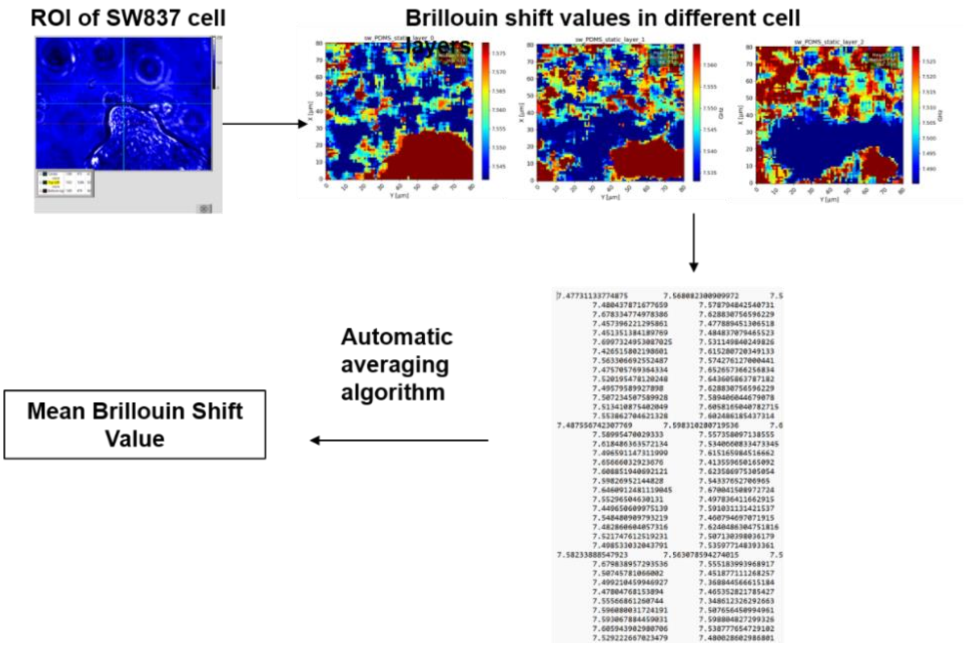


**Figure S3:** Workflow illustrating Brillouin analysis. An automated segmentation algorithm identifies the cell region of interest (ROI) and averages Brillouin Shift values across all voxels within the cell volume, providing whole-cell viscoelastic properties. This approach does not discriminate between intracellular structures, while the segmentation effectively restricts counting voxels only within the cell boundary.

#### S4. Phalloidin actin filament angle distribution

HIEC-6 and SW837 cells on PDMS (as static controls or following 24hrs of peristalsis stimulation) were stained with fluorescently conjugated phalloidin to visualize cytoskeletal networks and actin alignment. To quantify actin filament angle orientation, green channels images were isolated in Fiji. A single-pixel-line filter was applied at sequential rotational intervals (0°, 1°, 2°, etc.) and the brightness of the remaining pixels were summed. Then, the summed brightness at each angle was computed into a fraction. The totality of the 180 fractions equaled 100%. Angles were then grouped into increments of 30° by adding the fractions from 1° to 30°, 30° to 60°, etc. to assess the angle distribution trends.

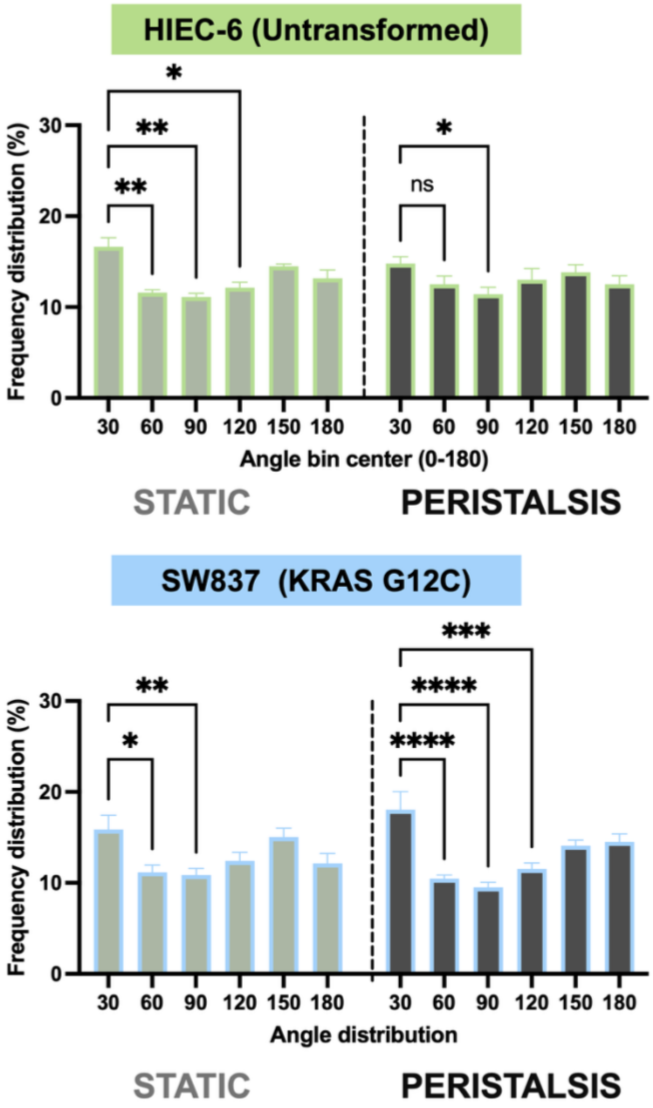


**Figure S4**. Frequency distribution of the phalloidin-stained actin filaments in HIEC-6 and SW837 G12C cells in static controls or following peristalsis. Minimal changes in actin fiber distribution are observed in HIEC-6 cells, while a significant re-arrangement of distribution is observed in SW837 cells in response to peristalsis. (*$p<0.05$, **$p<0.01$, ***$p<0.001$, ****$p<0.0001$, two-way ANOVA, $n\geq3$).

### S.5 RAS inhibition via sotorasib

To determine sotorasib concentrations to maximize viability, an MTS assay (Abcam,Cambridge, UK) was used to assess viability of cells over time. SW837 KRAS G12C were treated with different doses of RAS inhibitor (sotorasib) ranging from 1 nM to 10 μM in 2D for 48 hours, following which absorbance was measured at 490 nm with the Cytation 7 microplate reader (BioTek, Winooski, VT). Viability of the cells were normalized to untreated controls and expressed as percentage. A total of at least 4 replicates for each n=3 replicates was used for both control (nontreated) and drug-treated cells. GraphPad Prism 9 was used to fit a 4-parameter sigmoidal dose-dependent response curve to the raw viability data and measure IC50 concentrations for the drug in G12C (A). Once an optimal dose was determined (1.04μM), inhibition of RAS effector activity was evaluating via western blots for levels of phosphorylated ERK, normalized to total ERK protein expression (B).

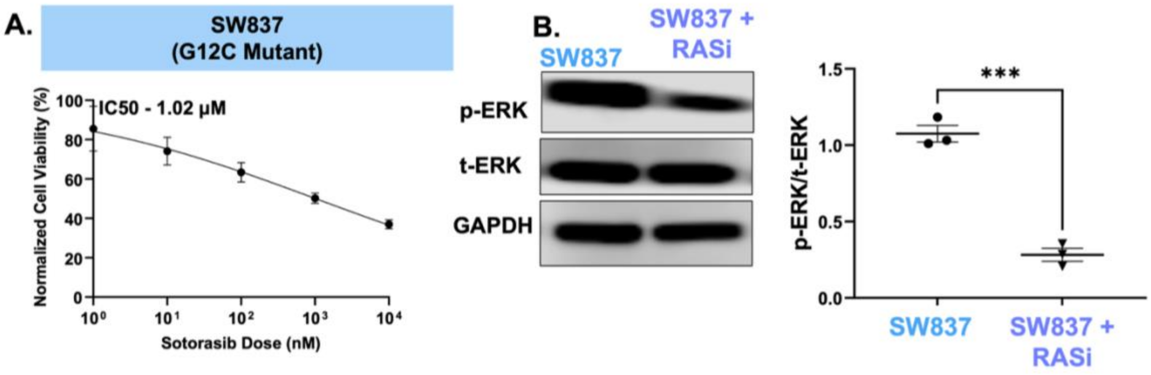


**Figure S5**. (A) Dose response curve for RAS inhibitor (sotorasib) against (A) HIEC-6 cells (B) SW837. Half-maximum inhibitor concentration (IC50) computed for cellular responses to drug compounds. (B) ERK activity in SW837 with 1.02 μM RASi (sotorasib) treatment. Representative western blot for p-ERK activity in SW837 cells after RASi treatment for 48 hours or maintained as controls. Band intensities were normalized to t-ERK and GAPDH as loading control. ***$p<0.001$, t-test, $n \geq 3$.

### S6. Effect of RAS inhibition (sotorasib) on baseline expression of LGR5 in static SW837 KRAS G12C cells

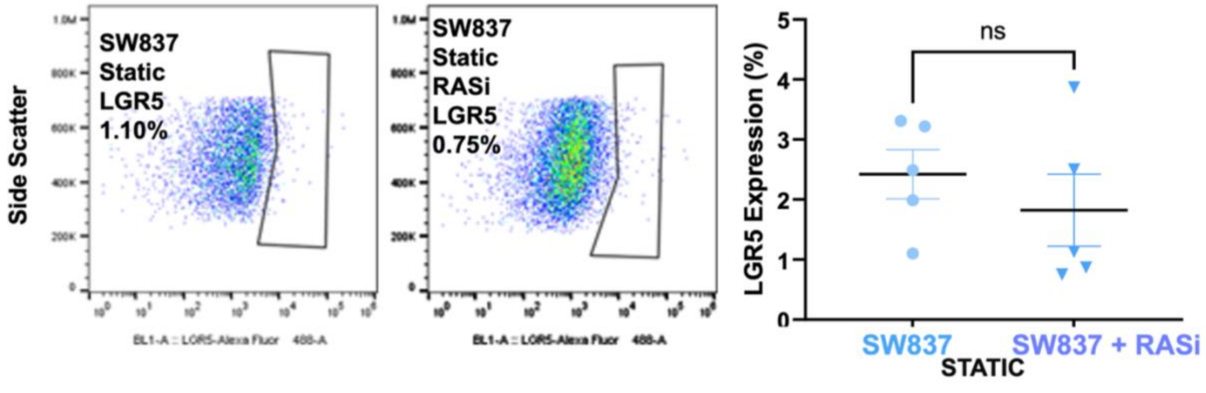

**Figure S6.** Representative LGR5 flow cytometry plots of SW837 cells maintained in static controls or treated with RASi. Individual dot plots summarizing flow analysis of SW837 KRAS G12C LGR5$^{+}$ expression (%) after 24hr exposure to RASi or maintenance in static controls. No change in LGR5$^{+}$ expression was noted in KRAS mutant cells exposed to RASi alone compared to static controls. (ns, t-test, n≥3).

**S.7 RAS expression in HIEC-6 G12C transfected cells**

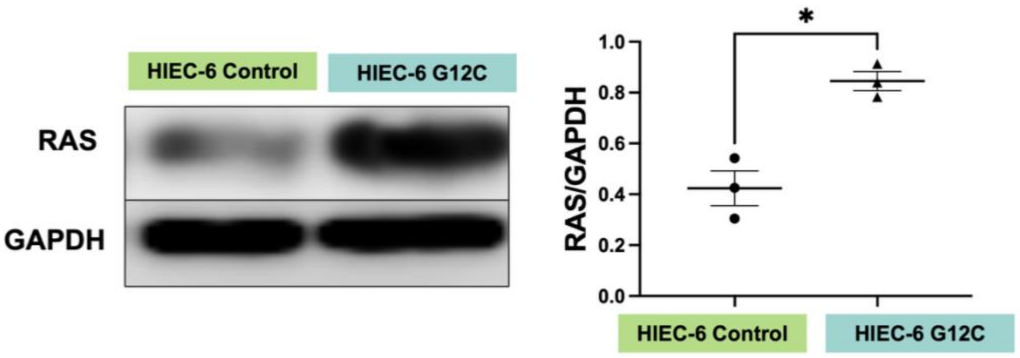


**Figure S7**. Representative western blots of RAS activity in KRAS G12C HIEC-6 cells compared to HIEC-6 control 2D cells after 72 hours post KRAS G12C transfection. Densitometric quantification of RAS activity normalized to corresponding GAPDH loading controls.(*$p<0.05$, t-test, n≥3) .

**S.8 Effect of KRAS G12C transfection on static HIEC-6 cultures (untrasformed or transfected)**

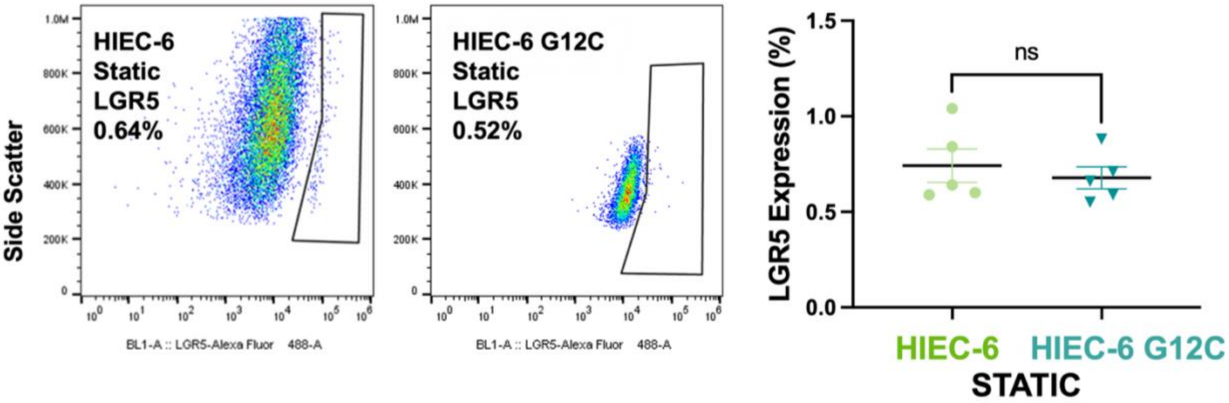


**Figure S8.** Representative LGR5 flow cytometry plots in KRAS G12C HIEC-6 cells compared to HIEC-6 control 2D cells after 48 hours post KRAS G12C transfection. No change in LGR5$^{+}$ expression was noted in KRAS G12C HIEC-6 cells alone compared to 2D controls.

### S.9 Flow Cytometry Gating Technique

Isotype controls were used to define a 0.5% background threshold, and polygon gates were drawn in FlowJo (Ashland, OR) to exclude this background signal. These gates were then applied unchanged to the antibody-stained samples to identify the positive population.

**For Figure 1A**

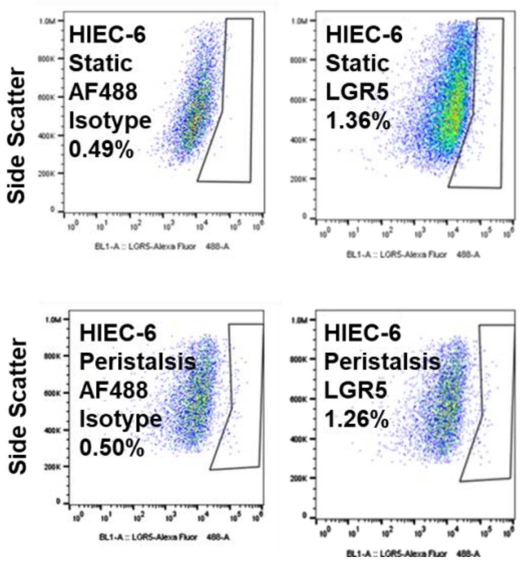


**For Figure 1B**

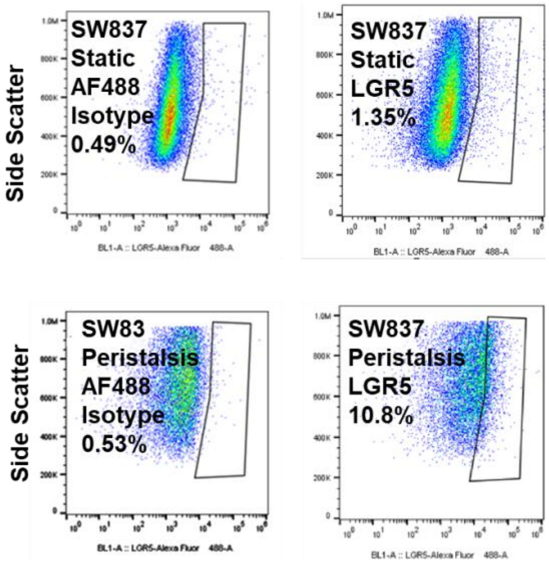


**For Figure 5C**

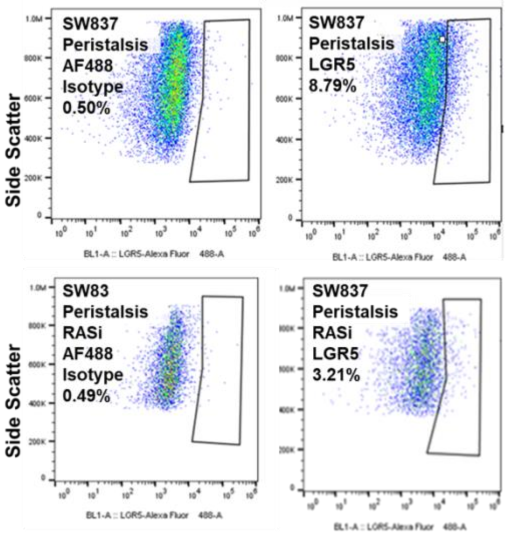


**For Figure 6C**

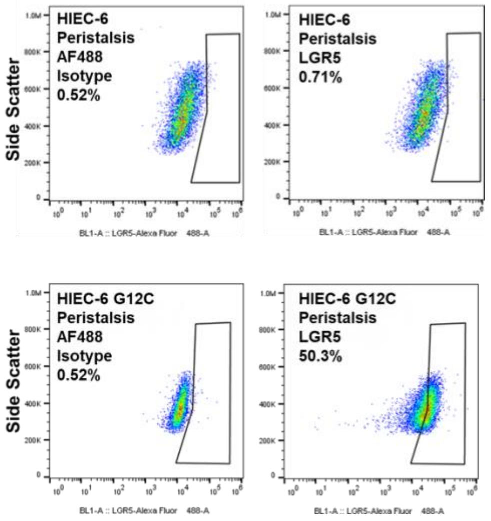


**Figure S9:** LGR5 flow cytometry gating technique using both unstained and isotype controls. Polygon gates (outlined in black) were drawn in FlowJo to define a 0.5% background cutoff on the isotype control plots (left). This gate was then applied directly to the antibody-stained samples (right) to quantify the percentage of cells expressing LGR5.

#### S.10 Manual quantification of Aspect Ratio

Actin filaments were visualized via green fluorescence and nuclei were visualized via blue fluorescence. With the use of Image J, calibration was set according to the image's scale bar. Major filament and minor filament were marked with the "straight line" functions in Image J. An example of the lines drawn is depicted in **Supplementary Figure S2.** In the "Set Measurements" feature, "length" was selected to track major and minor filament length. After each line was drawn, the "Measure" (CTRL+M) analysis tool was selected to produce a length value. Once all lines were drawn, the number of lines was extrapolated from the results box. Aspect Ratio was calculated as

$$\frac{Average\ Major\ Filament\ Length}{Average\ Minor\ Filament\ Length}$$

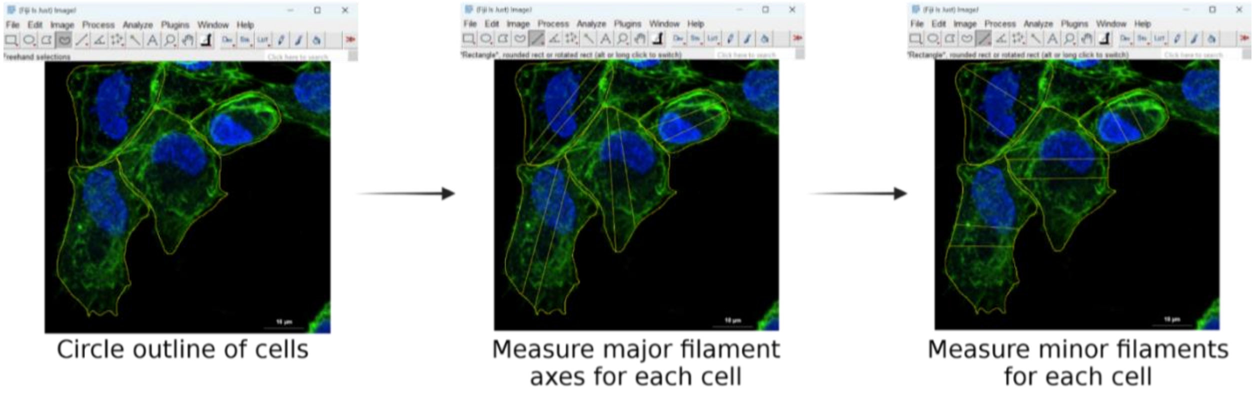


**Figure S11:** Representative images illustrating manual measurements of major and minor filament lengths used for aspect ratio analysis. A minimum of 4 ROIs were drawn per cell (two major and two minor filaments), and averaged values were used to calculate aspect ratio. Scale bar 10µm.

**S.11 Uncropped blots used for this study**

**For Figure 1C.**

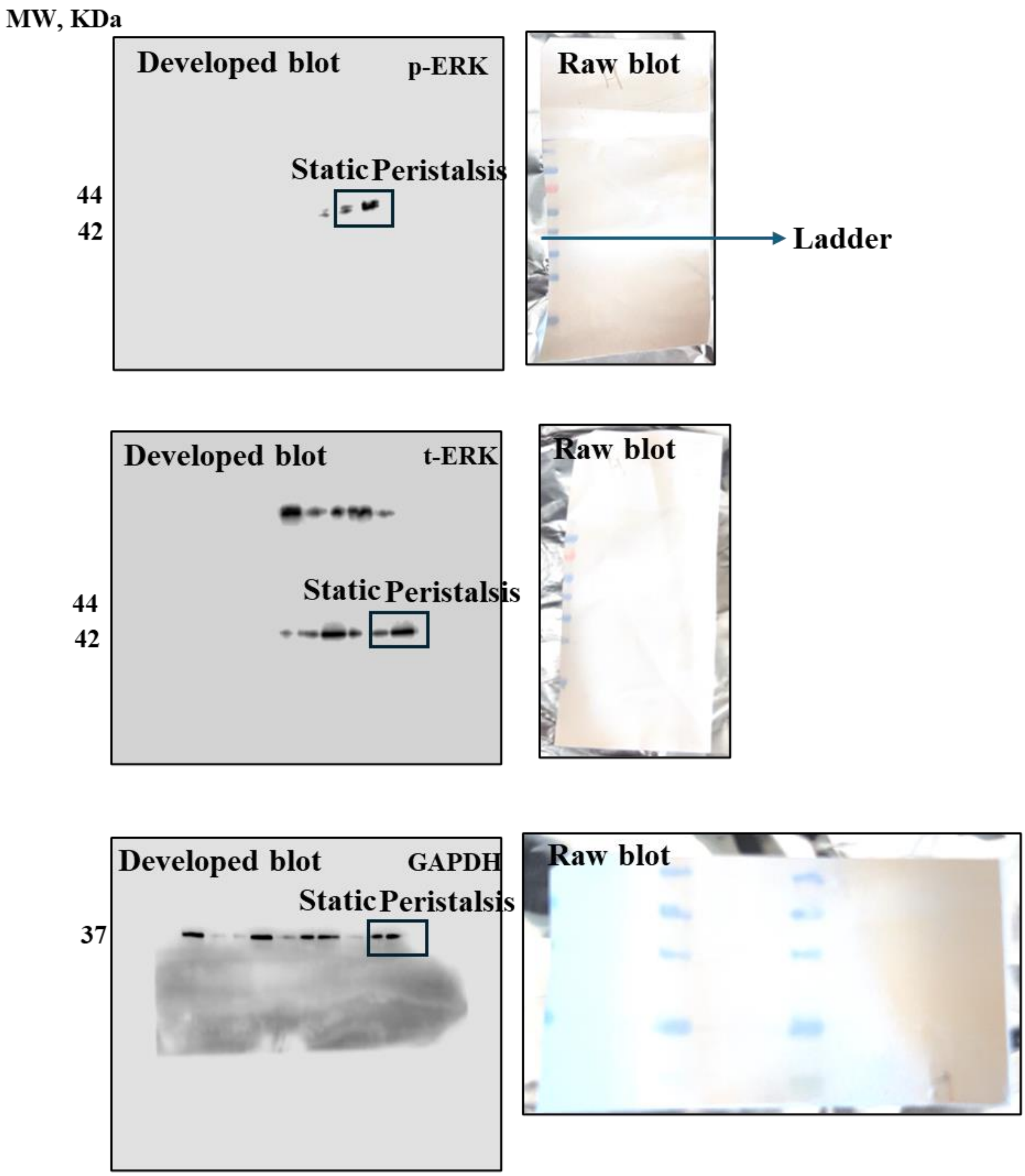

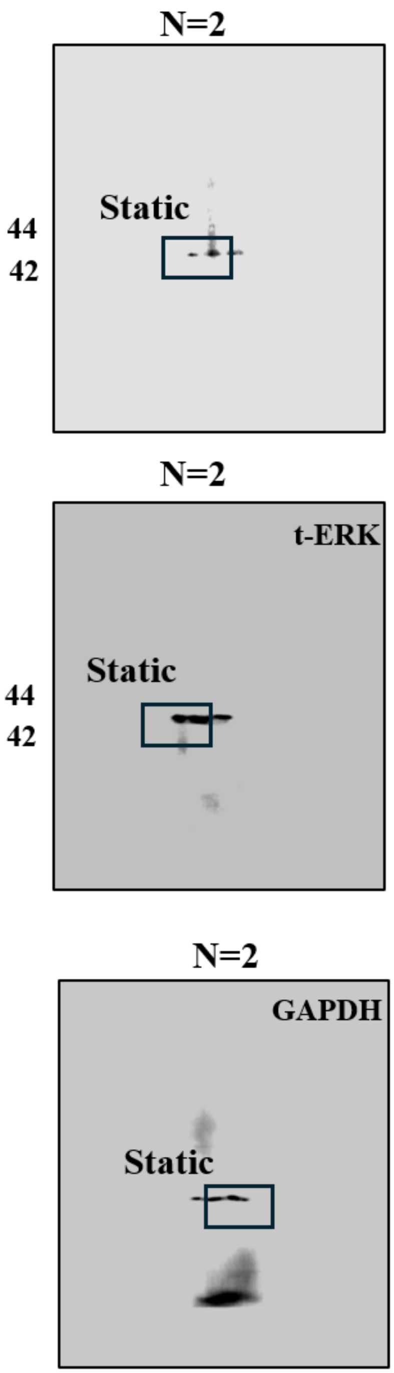
N=2
Static
44
42
N=2
t-ERK
Static
44
42
N=2
GAPDH
Static

**For Figure 1D**

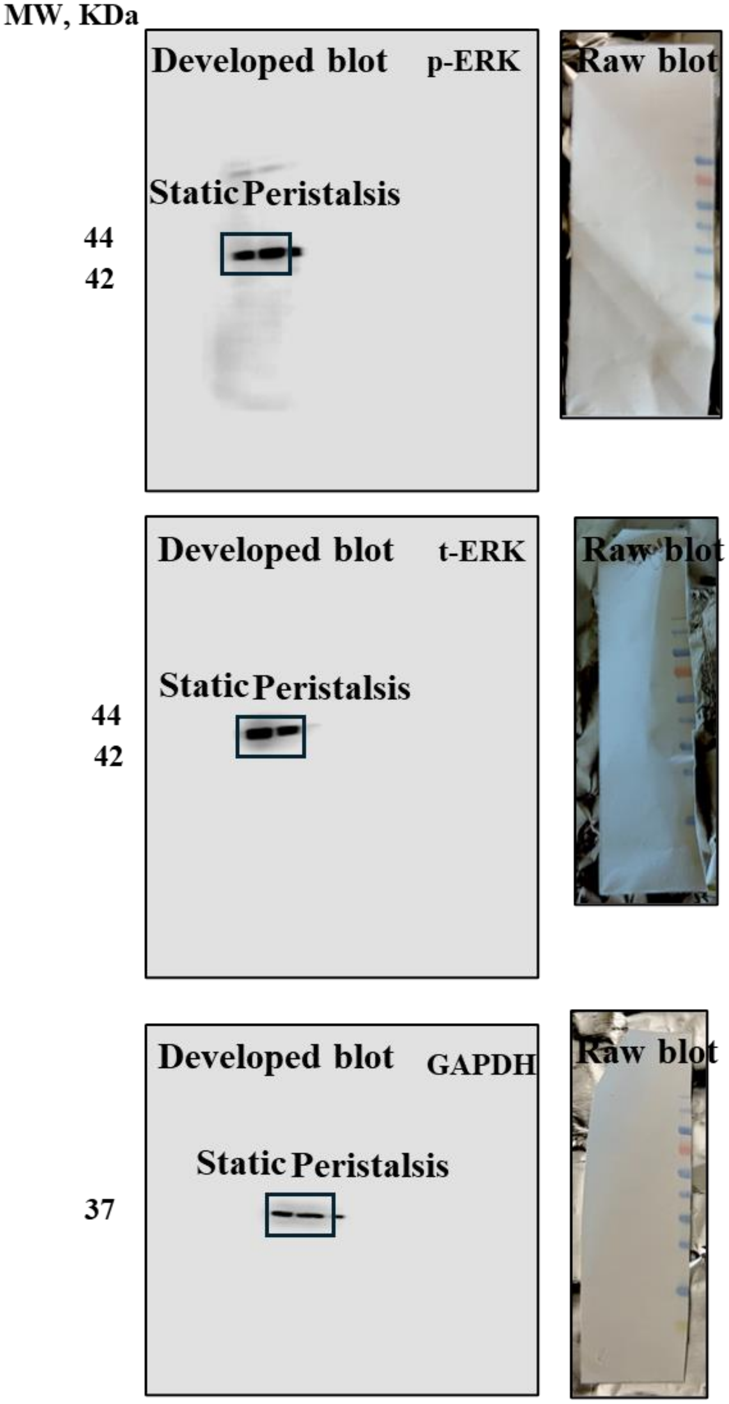

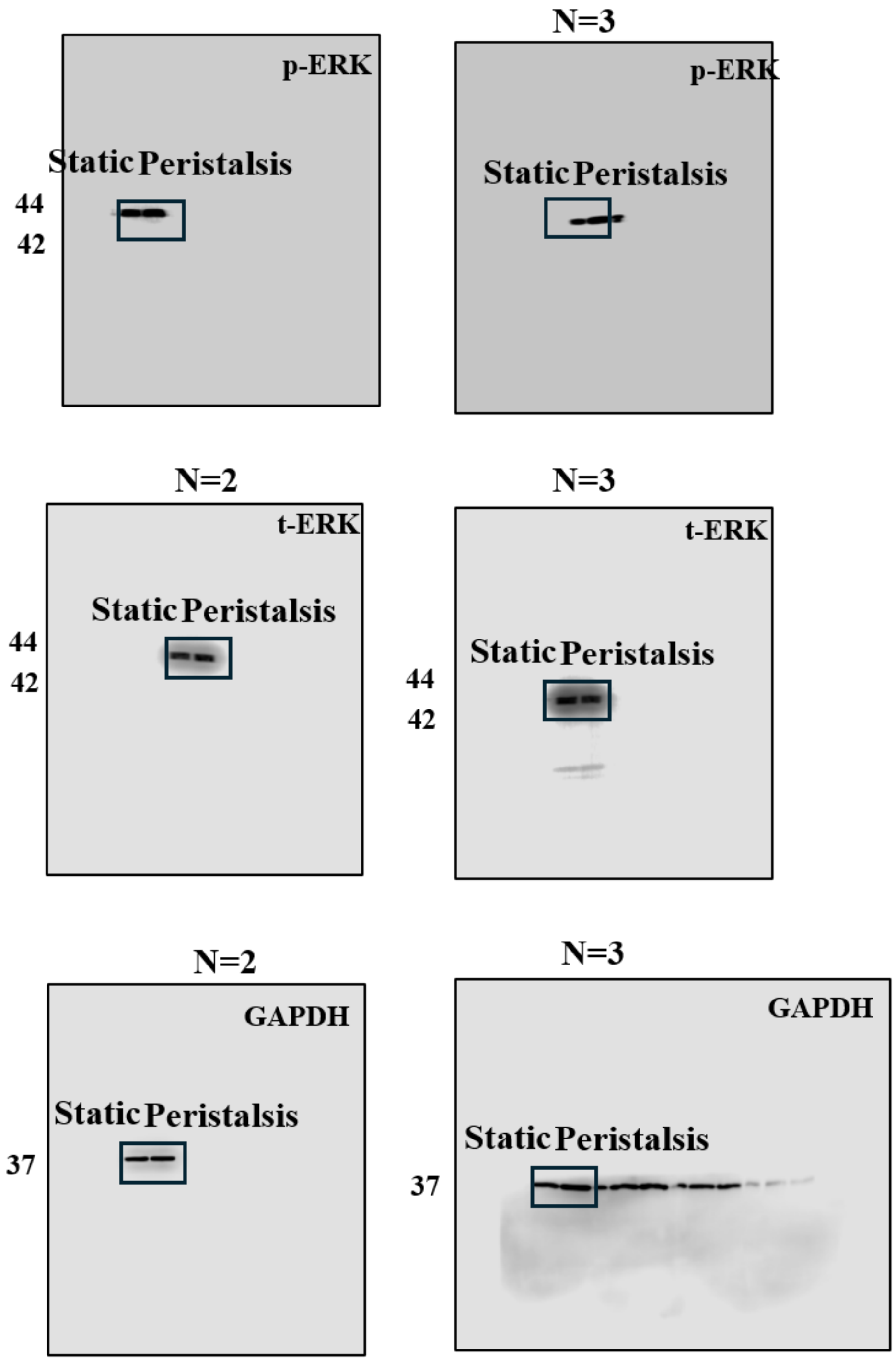
N=3
p-ERK
Static
Peristalsis
44
42
p-ERK
Static
Peristalsis
N=2
N=3
t-ERK
Static
Peristalsis
44
42
t-ERK
Static
Peristalsis
44
42
N=2
N=3
GAPDH
Static
Peristalsis
37
GAPDH
Static
Peristalsis
37

**For Figure S5B**

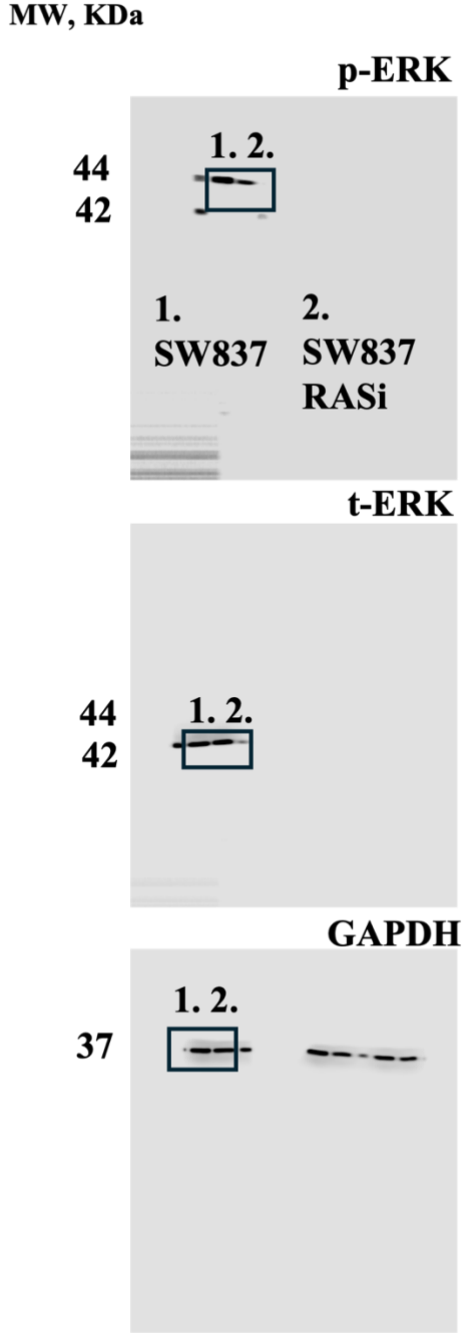


**For Figure S7**

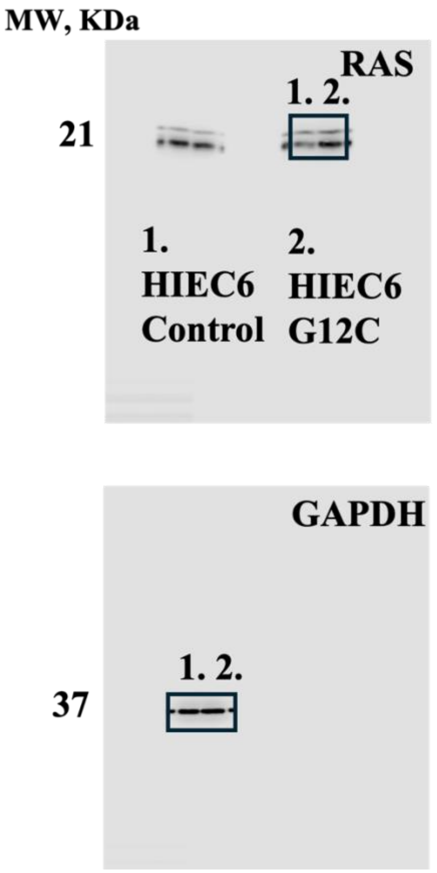


**Figure S11:** Uncropped western blots showing the boxed samples used in figures in the main text, with the protein assayed, it's molecular weight and loading controls. Raw blots for each representative blot are included, where appropriate which includes visible ladder bands and the membrane edges. The original membrane was sectioned prior to antibody hybridization; therefore, full length images intact membrane edges cannot be provided. All blots were processed in parallel, and only representative blots are shown.
